# An Algorithmic framework for genome-wide identification of Sugarcane (*Saccharum officinarum* L.)-encoded microRNA targets against SCBV

**DOI:** 10.1101/2020.10.25.353821

**Authors:** Muhammad Aleem Ashraf, Xiaoyan Feng, Xiaowen Hu, Fakiha Ashraf, Linbo Shen, Shuzhen Zhang

**Author notes:** Correspondence (M.A.A), (S.Z). These authors contributed equally to this work (Muhammad Aleem Ashraf, Xiaoyan Feng, Xiaowen Hu).

## Abstract

*Sugarcane Bacilliform Virus* (SCBV) is considered an economically the most damaging pathogen for sugarcane production worldwide. Three ORFs are characterized in a single molecule of circular, ds-DNA genome of the SCBV, encoding for hypothetical protein (ORF1), DNA binding protein (ORF2) and Polyprotein (ORF3). The study was aimed to predict and comprehensively evaluate sugarcane miRNAs for the silencing of SCBV genome using *in-silico* algorithms. Computational methods were used for prediction of candidate miRNAs from sugarcane (*S. officinarum* L.) to silence the expression of SCBV genes through translational inhibition by mRNA cleavage. Mature sugarcane miRNAs were retrieved and were assessed to hybridization with the SCBV genome. A total of fourteen potential candidate miRNAs from sugarcane were computed by all the algorithms used for the silencing of SCBV. A consensus of three algorithms predicts hybridization sites of sof-miR159e at common locus 5534. The miRNA-mRNA interaction was estimated by computing free-energy of miRNA-mRNA duplex using RNAcofold algorithm. Regulatory network of predicted candidate miRNAs of sugarcane with SCBV ORFs, generated using Circos, identify novel targets. Consequently, detecting and discarding inefficient amiRNAs prior to cloning would help suppressed mutants faster. The efficacy of predicted candidate miRNAs was evaluated to test the survival rate of the *in vitro* amiRNA-mediated effective badnaviral silencing and resistance in sugarcane cultivars.

## 1. Introduction

Sugarcane bacilliform viruses (SCBVs) are classified into Badnavirus genus of Caulimoviridae family. These viruses are composed of monopartite, circular, non-enveloped bacilliform with dimensions (30nm ×120-150 nm), double-stranded DNA (ds-DNA)-genome of approximately 7.2-9.2 Kbp in size [1]. The genome of SCBV constitutes three major open reading frames (ORFs) that are located on the ‘plus DNA strand’ with a single discontinuity [2]. ORF1 encodes a small hypothetical protein which has unknown function. ORF2 encodes a virion-associated DNA-binding protein. ORF3 encodes a largest polyprotein which is represented as (P3.) that was composed of multiple functional sub-units. The polyprotein (P3) is cleaved by a viral aspartic protease to give rise multiple functional small proteins: intracellular movement, capsid, aspartic protease, RT, and ribonuclease [1–6]. H. RT-RNaseH-coding region was considered the most common taxonomic marker for identification of badnaviral genomic components. This coding region is a standard source to compare the sequence diversity of the badnaviral genomes [7]. The *Badnavirus* genome is replicated using reverse transcription of a more-than-genome length RNA. It works as a standard template to complete the translation process of the badnavirus proteins. It is also used as reverse transcription for replication of the badnavirus genome [8, 9].

First report of SCBV infection was observed in the Cuban sugarcane cultivar B34104 in 1985 [10].These viruses are disseminated worldwide and decrease crop production significantly because of accessibility and exchange of biological materials globally. SCBV is a source of infection for several bioenergy crop sugarcane cultivars, varieties and species. Broad host range of SCB includes diverse and economically important members of *Poaceae* (sugarcane, rice) and *Musaceae* (banana) families. Natural transmission of SCBV is disseminated by sap-feeding mealy bug species by vegetative cutting [11]. SCBV disease symptoms include chlorosis and leaf freckle. Infected sugarcane plants were also monitored to have no symptoms. SCBV-infection in sugarcane plants causes much reduced sucrose content, juice, stalk weigh, purity and gravity in recent years in China [6].

Plant employs multiple defense strategies to control virus invasion, replication and transmission in the cell. RNAi-mediated response of plants against invading viruses is highly significant during infection period [12]. RNAi mechanism inhibits the protein translation at mRNA level using a highly sequence specific strategy [13]. Sugarcane plant species were imparted natural inherited active immunity consisting of small, non-coding microRNAs (miRNAs) to control viral diseases. The miRNA-mediated genome silencing was considered to certify the activity of positive or negative immune-based regulation. It is also considered as key activator of immune defense in plants [14]. These miRNAs belong to a class of 21-23 nucleotides long, endogenous, non-coding RNA molecules that plays a major role to control the gene expression at post-transactional level. RNA silencing in the form of miRNAs within the host plant, is a source of natural immunity. It provides resistance to the host plant after infection of foreign genetic elements including plant viruses [15–17]. Artificial microRNA (amiRNA)-mediated silencing of invading viruses in plants was first time reported by Niu [18]. This amiRNA-based silencing strategy was applied effectively in many plants to combat plant viruses: cotton leaf curl Kokhran virus (CLCuKoV-Bu) [19], Cucumber mosaic virus (CMV)[20], CymMV and ORSV [21]. Serological method, ELISA, immunosorbent electron microscopy (ISEM) [22], IC-PCR [23], RT-PCR [24], rolling circle amplification (RCA) [25] are the most widely used reliable detection techniques for SCBV. New strategies are under development for improved diagnoses and cure of badnaviruses [26].

Prediction and validation of host-virus interactions is the basic step to understand the critical functional efficacy of miRNAs in the broader range of miRNA-mRNA regulatory networks controlling biological processes. Each target miRNA possesses potential hybridization binding sites. In order to validate potential miRNA target experimentally is highly costly and laborious. An *In Silico* approach to identify miRNA targets concludes potential target binding sites for laboratory research. Computational strategies determine how miRNAs target to a desirable mRNA. A large number of computational algorithms are publically available for miRNA target prediction. It is highly advantageous to get biological access to several computational tools with different features. The researcher is challenged with an important choice for selecting suitable tools for prediction [27]. The current study was designed to implement miRNA prediction algorithms and identify potential targets of sugarcane-derived miRNAs against SCBV as precedent for creating resistance to sugarcane cultivars using RNAi technology. We have also screened the potential sugarcane miRNAs for understanding sugarcane-badnavirus interactions. The novel amiRNA-mediated approach is credible to generate SCBV-resistance sugarcane plants through genetic engineering.

## 2. Materials and Methods

### 2.1. Retrieval of Sugarcane MicroRNAs

The sugarcane mature microRNAs (miRNAs) and stem-loop hairpin precursor sequences were retrieved from the miRNA biological sequence database, miRBase (v22) (http://mirbase.org/). miRBase serve as primary public repository and standard online reference resource for all published miRNA sequences, textual annotation and gene nomenclature [28–30]. In this study, 16 *S.officinarum* (MI0001756-MI0001769) and 19 Saccharum spp. (MI0018180-MI18197) miRNA sequences were downloaded (Table S1).

### 2.2. SCBV Genome Retrieval and Annotation

The full-length transcript of the SCBV-BRU genome (Genbank accession number JN377537.1, size 7884 bp) was isolated from the S.officinarum cultivar, cloned, sequenced and submitted to the NCBI database [31]. Expected size and abundance of ORFs along nucleotide distribution of the above mentioned NCBI retrieved SCBV-BRU genome was estimated using pDRAW32 DNA analysis software (v 1.1.129). The SCBV-BRU genome annotation is designed which represents ORFs of varying lengths.

### 2.3. Target Prediction in SCBV Genome

Target prediction is considered a key feature towards identification of credible miRNA-mRNA interactions hybridization. A present, many target prediction algorithms was designed to predict and identify best candidate miRNA targets. Each tool uses a specific criteria and stringency for miRNA target prediction. We used four target prediction algorithms cited in literature — miRanda, RNA22, RNAhybrid and psRNATarget to find the most relevant sugarcane miRNAs for silencing of SCBV genome (Table 1). We designed the most effective computational approach to analyze miRNA targets at three different prediction levels: individual, union and intersection. A detailed workflow pipeline was present in (Fig 1).

**Fig 1.**
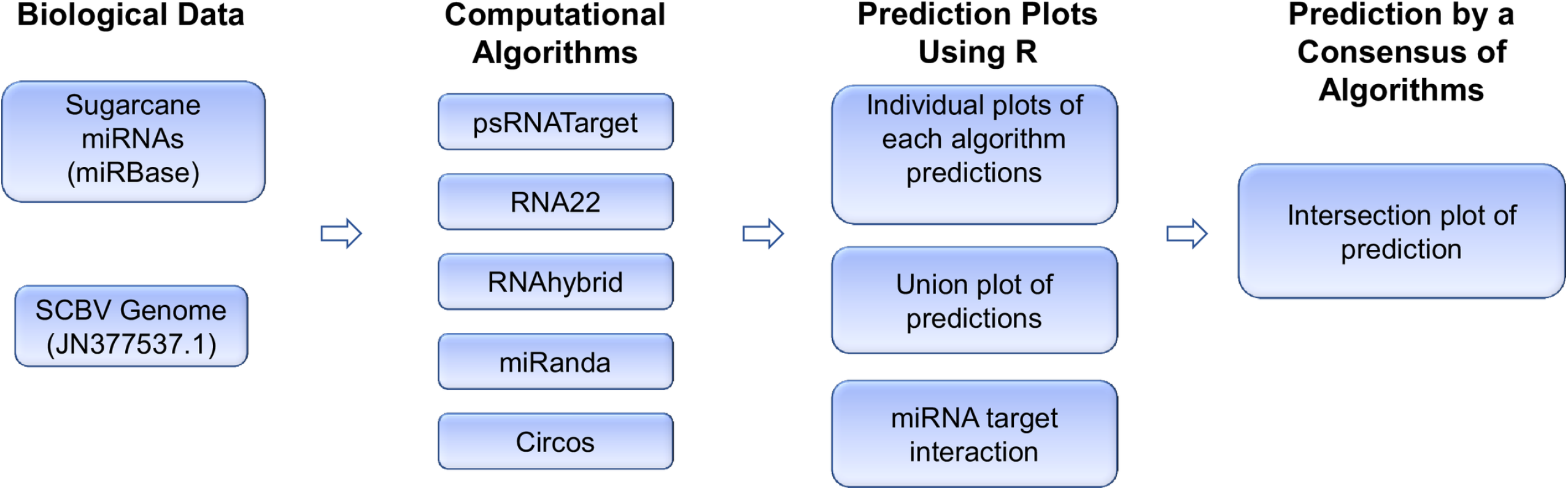
A flowchart embodies the methodology of miRNA prediction from the SCBV genome. A flowchart designed for predicting candidate miRNAs from the SCBV genome pipeline. Biological data composed of sugarcane miRNAs retrieved from the miRBase database and SCBV genome from NCBI genbank database. An algorithmic framework consists of three kinds of tools used for identification of sugarcane-encoded miRNA targets, prediction of secondary structures and visualization of miRNA-target interaction. R language was used to make plots and select data using in-house scripts/codes.

**Table 1.** Comparison of distinctive parameters used in the common target prediction tools.

| Tools | Algorithms | Seed pairing | Target site accessibility | Multiple sites | Translation Inhibition | Availability |
| --- | --- | --- | --- | --- | --- | --- |
| miRanda | Local alignment | + | + | + | + | Web server and source code |
| RNA22 | FASTA | – | + | + | – | Only web server |
| RNAhybrid | Interamolecular hybridization | + | + | + | + | Web server and source code |
| psRNATarget | Smith-Waterman | – | + | + | + | Only web server |
| Tapirhybrid | FASTA | + | + | + | – | Web server and source code |
| Targetfinder | FASTA | + | – | – | – | Only source code |
| Target-align | Smith-Waterman | – | – | + | – | Web server and source code |
| Targetscan | Custom made | + | – | + | + | Only source code |

‘+’ Represent feature used, ‘-’ indicates that these features were not used.

### 2.4. miRanda

miRanda is considered more frequently used standard miRNA-target predictor scanning algorithm. It was implemented first time in 2003 [32] and updated into a web-based tool for miRNA analysis [33]. It considers following algorithmic features; seed-based interaction, minimum free energy (MFE), sequence complementarity and interspecies conservation in the target genome. The latest version of miRanda software was accessed using online source website (http://www.microrna.org/). SCBV genome and sugarcane mature miRNA sequences were processed after defining default setting. These include energy threshold = −15kcal/mol, score threshold = 130, scaling parameter = 4.00, gap-open penalty= −9.00 and gap-extend penalty= −4.00.

### 2.5. RNA22

RNA22 is a friendly user, highly sensitive, web-based (http://cm.jefferson.edu/rna22v1.0/) novel pattern-recognition algorithm that serves as interactive exploration for predicting target sites with corresponding hetero-duplexes. Non-seed based interaction, pattern recognition, site complementarity and folding energy are key parameters of RNA22 algorithm [34]. Final scoring obviates to use cross-species conservation sequence filter [35]. The minimum number of paired-up bases in heteroduplex, maximum folding energy for heteroduplex and maximum number of G: U wobbles allowed in the seed region are user-defined parameters. RNA22 was processed under default parameters: specificity (61%), sensitivity (63%), and seed size (7) with unpaired base (1), no limit for G: U wobbles, paired-up bases (12) and folding energy (−12.00 kcal/mol).

### 2.6. RNAhybrid

RNAhybrid is easy, fast, flexible web-based (http://bibiserv.techfak.uni-bielefeld.de/rnahybrid) intermolecular hybridization algorithm used to estimate miRNA-mRNA interaction as well target prediction based on MFE hybridization. The major parameters include MFE (energy threshold), target-site abundance and site complementarity. A p-value is assigned to assess RNA-RNA interaction based hybridization sites in the 3’ UTR sequence [36]. RNAhybrid is widely used to estimate the MFE of the consensus miRNA-target pair and the mode of target inhibition as suggested [37]. MFE threshold (−20Kcal/mol), hit per target (1), no G: U binding in seed region is fixed default parameters.

### 2.7. psRNATarget

psRNATarget is a newly developed web server (http://plantgrn.noble.org/psRNATarget/) to identify target genes of the plant miRNAs based on complementary matching scoring schema. It was used to discover validated miRNA-mRNA interactions [38]. The plant psRNATarget was designed to integrate key function for miRNA target prediction using complementarity scoring and secondary-structure prediction [39]. Target-site accessibility was evaluated by estimating unpaired energy (UPE) to unfold secondary structure [37]. We consider following unique parameter for analysis: (Expectation= 8.5, penalty for extending gap = 0.5, penalty opening gap = 2.0, penalty for G.U pair = 0.5) The HSP size = 19 and seed region = 2–13 nucleotides were set. Translation inhibition range was set 10-11 nucleotides.

### 2.8. Mapping of miRNA-Target Interaction

Interaction map was created between sugarcane miRNAs and SCBV ORFs using the Circos algorithm [40].

### 2.9. RNAfold

RNAfold is a new web-based algorithm applied for the prediction of stable secondary structure of pre-miRNAs based on the MFE [41].

### 2.10. Free energy (ΔG) Estimation of Duplex Binding

RNAcofold is a novel web-based server applied for the estimation of free energy (ΔG) associated with miRNA-mRNA interactions [42]. Free energy of miRNA: miRNA duplex is considered a key predictor for miRNA targets during hybridization. FASTA sequences of sugarcane consensus miRNAs and their corresponding targets were processed to RNAcofold using default parameters.

#### 2.10.1. Graphical Representation

All the computational data was processed into graphical representations using R language (v3.1.1) [43].

## 3. Results

### 3.1. Genome Assembly of SCBV

SCBV is a plant pararetrovirus, was classified in the genus *Badnavirus* of the family *Caulimoviridae*. The genomic ds-DNA molecule of SCBV is comprised of three ORFs, separated by an intergenic region (IR). ORF1 is composed of 557 nucleotides (618-1175 nt) encoding a hypothetical protein (P1) with 185 amino acids (aa). While ORF2 composed of 370 nucleotides (1176-1546 nt) codes for a virion-associated DNA binding protein (P2) with 123 aa. The precise functional capabilities of these proteins (encoded by ORF1 and ORF2) have not been explored. A large polyprotein (1977 amino acids) is encoded by ORF3 (1547-7479 nt) to cleave by a viral aspartic protease. The resulting proteins obtained are named as movement, capsid protein, aspartyl proteinase, reverse transcriptase and ribonuclease H. The IR is composed of 1022 nucleotides (7479-618) and is located between 3’-ORF3 to 5’-ORF1. The IR works as a promoter and controls the transcription and regulation of the SCBV genome. Genome organization of the SCBV with three ORFs is shown in (Fig 2).

**Fig 2.**
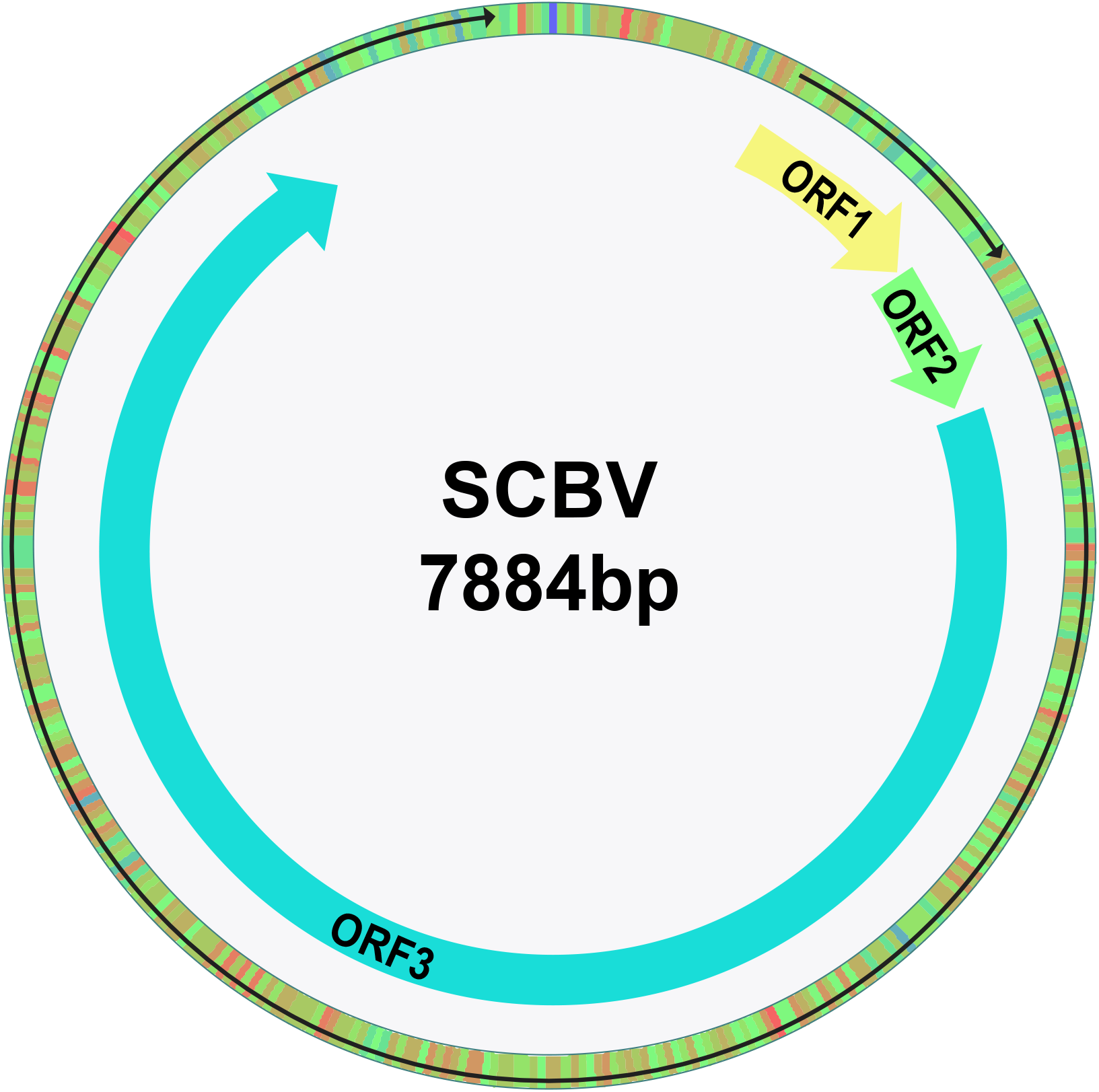
Genomic organization of the Sugarcane Bacilliform Virus. Genome organization of the Sugarcane Bacilliform Virus showing the predicted ORFs with arrows was composed dsDNA of 7884 bp in size.

### 3.2. ORF1 encoding Hypothetical Protein

The hypothetical protein of the SCBV genome was encoded by the ORF1 which has unknown function [2]. miRanda predicted only two target sites of sugarcane miRNAs (sof-miR156 and sof-miR168) at nucleotide positions (818-837 and 617-638) to target ORF1 (Fig 3A). RNA22 predicted binding sites of miRNAs (sof-miR156 and sof-miR168a) at two different locus positions 817 and 834 respectively (Fig 3B). RNAhybrid algorithm predicted multiple potential binding sites of sugarcane miRNAs (sof-miR168 (a, b), ssp-mi827, ssp-miR1128) at multiple nucleotide positions (612-632, 1170-1192, 1137-1157) respectively (Fig 3C). In addition, psRNATarget identified potential hybridization sites of sof-miR159(c, e) at locus positions 1003 and 820 respectively (Fig 3D).

**Fig 3.**
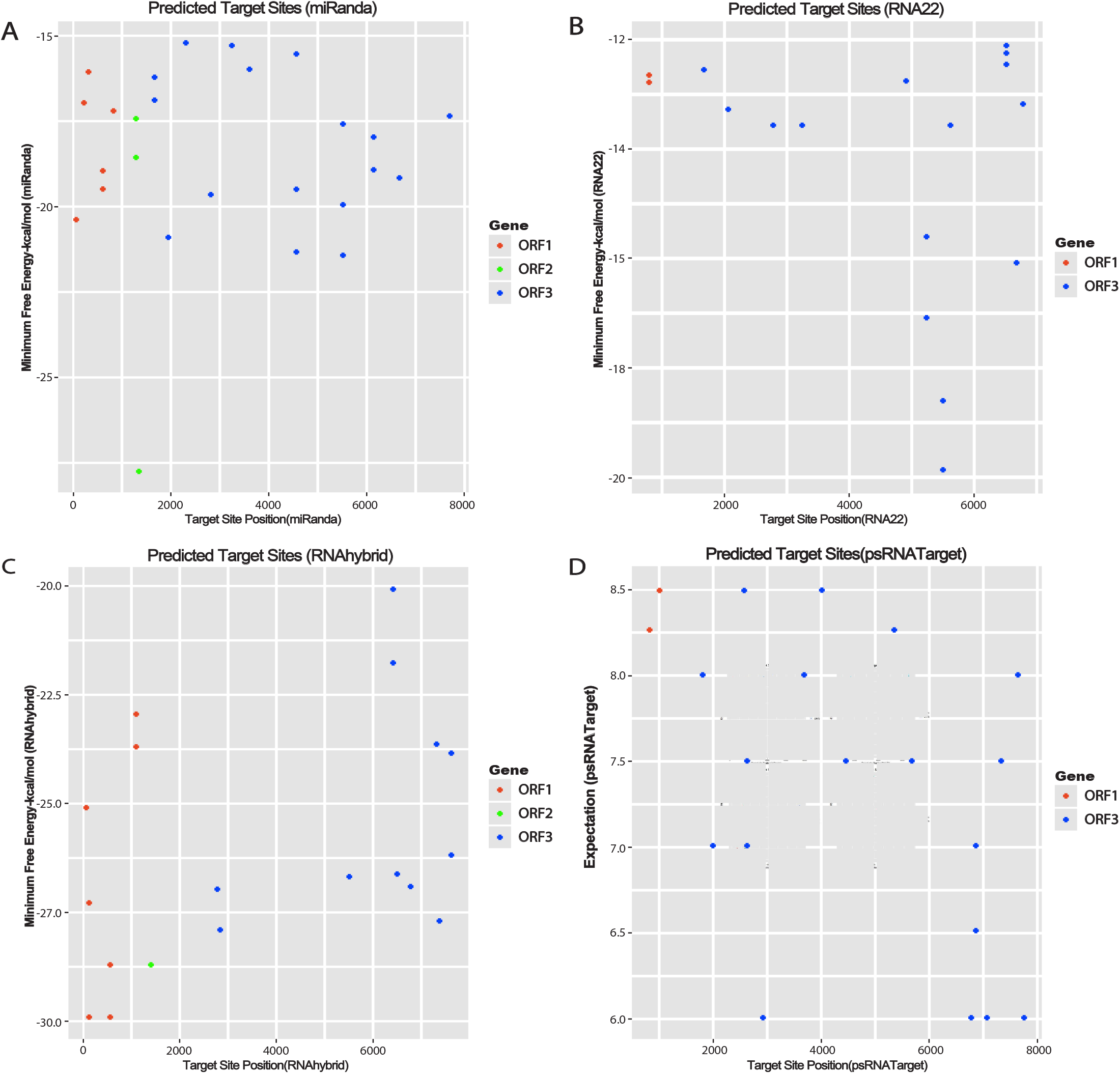
Target prediction of sugarcane miRNAs in the SCBV genome. Computational prediction of candidate miRNA targets in the genome of the SCBV. A) miRNA-targets obtained from miRanda. B) RNA22 predicted potential hybridization sites C) Target sites of sugarcane miRNAs as identified by RNAhybrid. D) Prediction results of target sites of sugarcane miRNAs obtained by psRNATarget.

### 3.3. ORF2 encoding DNA Binding Protein

Nucleic acid (DNA)-binding protein of the SCBV genome was encoded by ORF2 [6, 44]. RNAhybrid and miRanda predicted potential target binding site of ssp-miR166 at locus position (1449-1470) (Fig 3A and 3C). Suitable candidate miRNAs from sugarcane (ssp-miR444 (a, b, 3p) was observed to target the ORF2 at single loci nucleotide position (1301-1326) determined by miRanda algorithm (Fig 3A). No sugarcane miRNAs were predicted to target ORF2 gene by RNA22 tool (Fig. 3B). Similarly, RNAhybrid predicted bindings of ssp-miR166 at locus 1450 as shown in (Fig 3C). miRNA prediction results revealed that no candidate miRNA was identified to have potential genome-binding sites in the ORF2 region as predicted by psRNATarget as shown in (Fig 3D).

### 3.4. ORF3 encoding Polyprotein (CP, AP, RT and RNase H)

The poly proteins constitute the largest portion of the SCBV genome was encoded by the ORF3 [2, 6]. Potential candidate miRNAs from sugarcane were identified by miRanda algorithm to target the ORF3: (sof-miR159 (a, b, c, d, and e) at common locus 5534, sof-miR167 (a, b) at locus 2273, sof-miR168b at locus 4588, sof-miR408 (a, b, c, d, and e) at two common locus 4595 and 6695, ssp-miR166 at locus 1986, ssp-miR827 at locus 2816 and ssp-miR444 (a, b and c-3p) at common locus 6184. Multiple loci interactions were predicted for sof-miR159, sof-miR408 and ssp-miR444 families at nucleotide positions (5534–5552, 5576–5596), (4595–4615, 6695–6715) and (1679-1701, 3293-3313) of ORF3, respectively (Fig 3A).

Potential target binding sites were determined in the ORF3 of the SCBV genome by RNA22 algorithm: (sof-miR168a at locus 3263, sof-miR168b at nucleotide positions (1693, 3263), sof-miR396 at locus 2050 and ssp-827 at locus 2796 (Fig 3B). Multiple loci interactions were also identified for sof-miR159, sof-miR408 and ssp-miR444 families at nucleotide positions (5532, 6536), (5645, 6695) and (5246, 6793) respectively (Fig 3B).

Suitable miRNAs that potentially targeted the ORF3 were hybridized to understand miRNA-mRNA interaction, by RNAhybrid. These are sof-miR159 (a, b, d and e) at common locus 5535, sof-miR159c at locus 6518, sof-miR167 (a, b) at locus 2826, sof-miR169 at locus 7362, ssp-miR473 (a, b, c) at common locus 6438, ssp-miR444 (a, b) at locus 6796, ssp-miR444 c-3p at locus 2899 and ssp-miR1432 at locus 7314 (Fig 3C). ORF3 was targeted by candidate miRNAs: (sof-miR159e at locus 2647, sof-miR396 at locus 5363, ssp-miR166 at locus 1986, ssp-miR437 (a, c) at 2647 and ssp-miR827 at locus 7337 and ssp-miR444 (a, b and c-3p) at locus 6797) identified by psRNATarget. Multiple loci interactions were observed for sof-miR408 and ssp-miR444 families at nucleotide positions (1766-1786, 3669-3689, 5683-5702) and (4466–4486, 6797–6816, 6865-6885, 7079-7099) respectively (Fig 3D). Union plot indicates entire genome-binding sites identified by candidate miRNAs using target prediction tools (Fig 4, Table S2).

**Fig 4.**
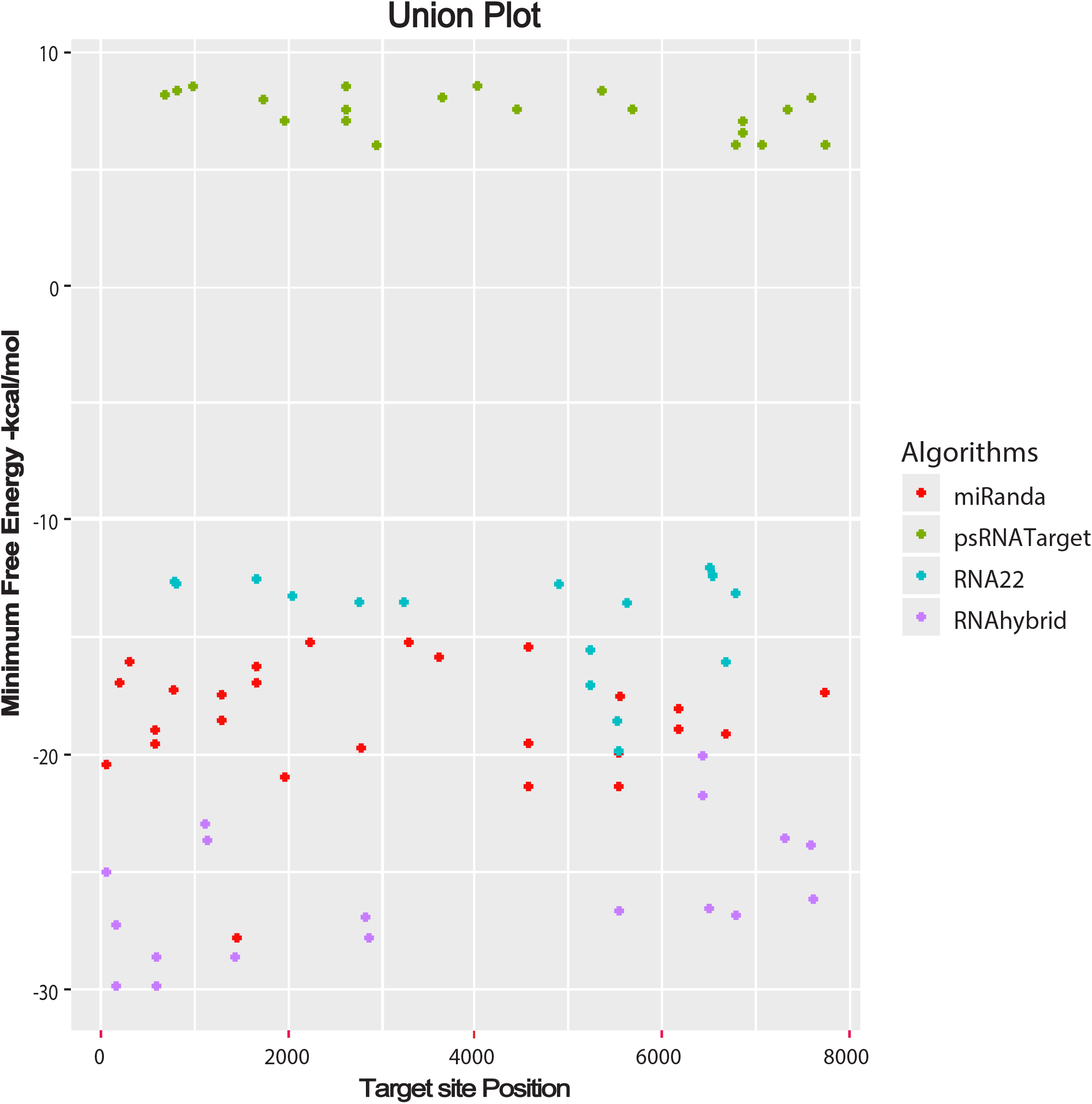
Union plot represents all the predicted sugarcane miRNA Targets in the SCBV genome. Union plot summarizes miRNA Target candidate prediction represented as a union from all the algorithms used in this study.

### 3.5. Visualization and Analysis of miRNA-Target Interaction Network

Though initially, Circos plotting tool was designed to analyse mutations in comparative metagenomics and transcriptomic biological data [45]. To study a comprehensive visualization of host-virus interaction, we created Circos plot to integrate biological data from sugarcane miRNAs and their predicted SCBV genomic target genes (ORFs) (Fig 5). In order to reduce visual graphical complexity and permit improved readability, we only used selected sugarcane miRNAs and their SCBV targets obtained from miRanda analysis. The results suggest that biological data visualization of candidate miRNAs from sugarcane, with SCBV-encoded ORFs determines credible information of desirable preferred targets of SCBV ORFs using consensus miRNAs. We have combined sugarcane miRNA data and their predicted SCBV targets simultaneously in this manner.

**Fig 5.**
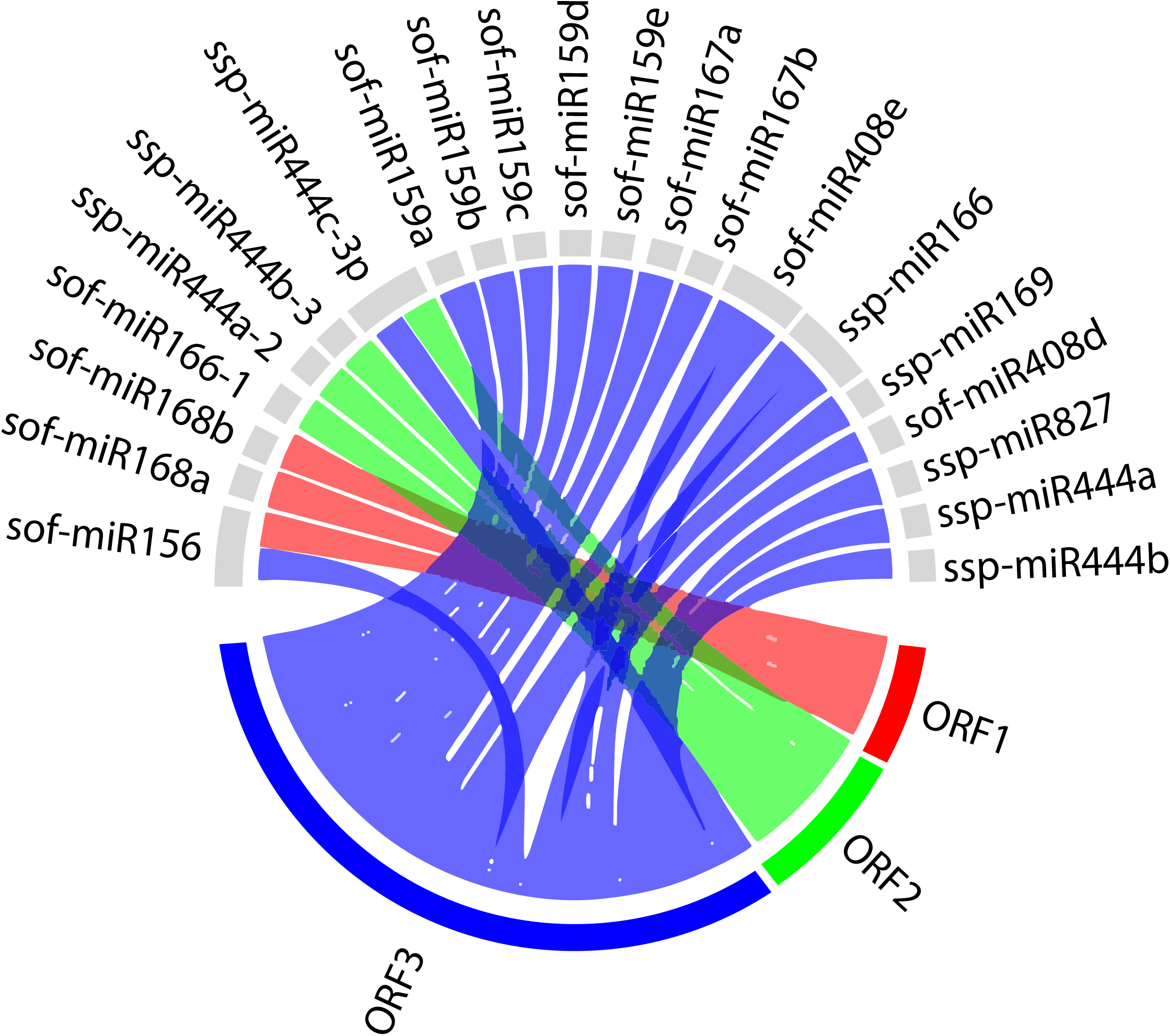
Circos plot representing miRNA-target interaction. Circos plot of the genomic regulatory network interaction predicted to be targeted by the sugarcane-miRNAs. The red, green and blue coloured lines represent SCBV genome components (ORFs). The synergetic counterparts of sugarcane miRNAs and their target genes (ORFs) of the SCBV genome are interconnected with coloured lines.

### 3.6. Predicting Common Sugarcane miRNAs

Based on predicted targeting miRNAs from sugarcane to silence the SCBV genome, fourteen miRNAs (sof-miR156, sof-miR159c, sof-miR159e, sof-miR168a, sof-miR396, sof-miR408a, sof-miR408b, sof-miR408c, sof-miR408d, sof-miR408e, ssp-miR827, ssp-miR444a, ssp-miR444b and sof-miR444c-3p) was detected by union of consensus multiple algorithms (miRanda, RNA22, RNAhybrid and psRNATarget) used in this study (Fig 6). Moreover, SCBV genomic components (ORF1, ORF2, ORF3 and LIR) were observed to be targeted by a total of eleven sugarcane miRNAs which were hybridized at unique positions within the ORF1: (sof-miR156 (locus 818), sof-miR168 (a, b) (locus 617) ORF2: ssp-miR166 (locus 1450) and ORF3: sof-miR159c (locus 5534), sof-miR408 (a, b, c, d and e) (locus 6695) and LIR: sof-miR396 (locus 79) according to intersection of two consensus algorithms (Table 2, Table S3).

**Fig 6.**
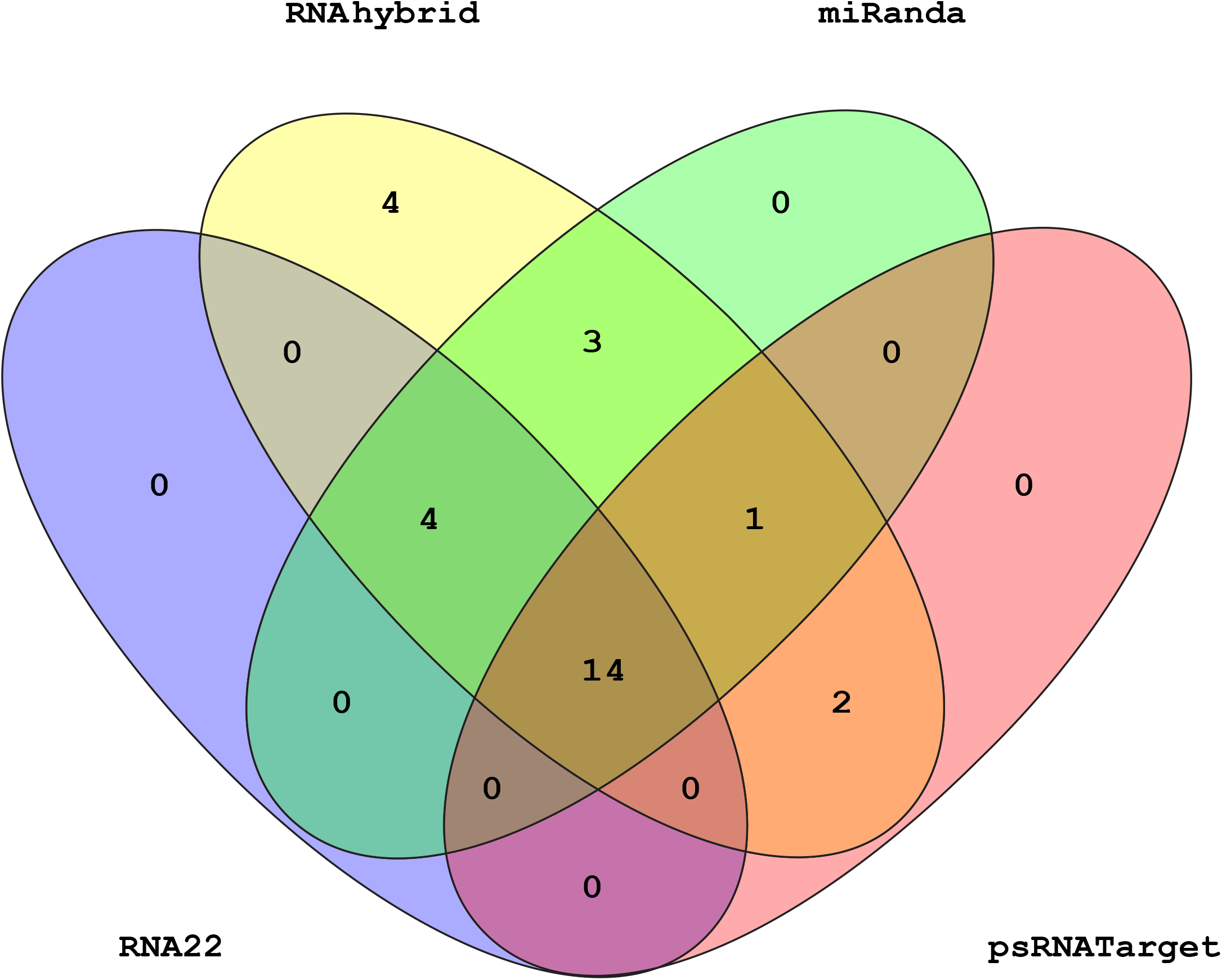
Venn diagram plot of SCBV genome targeted by sugarcane miRNAs. Venn diagram plot of SCBV genome targeted by sugarcane miRNAs. In total, 28 loci are targeted by sugarcane-miRNAs, as predicted from four unique algorithms.

**Table2:**
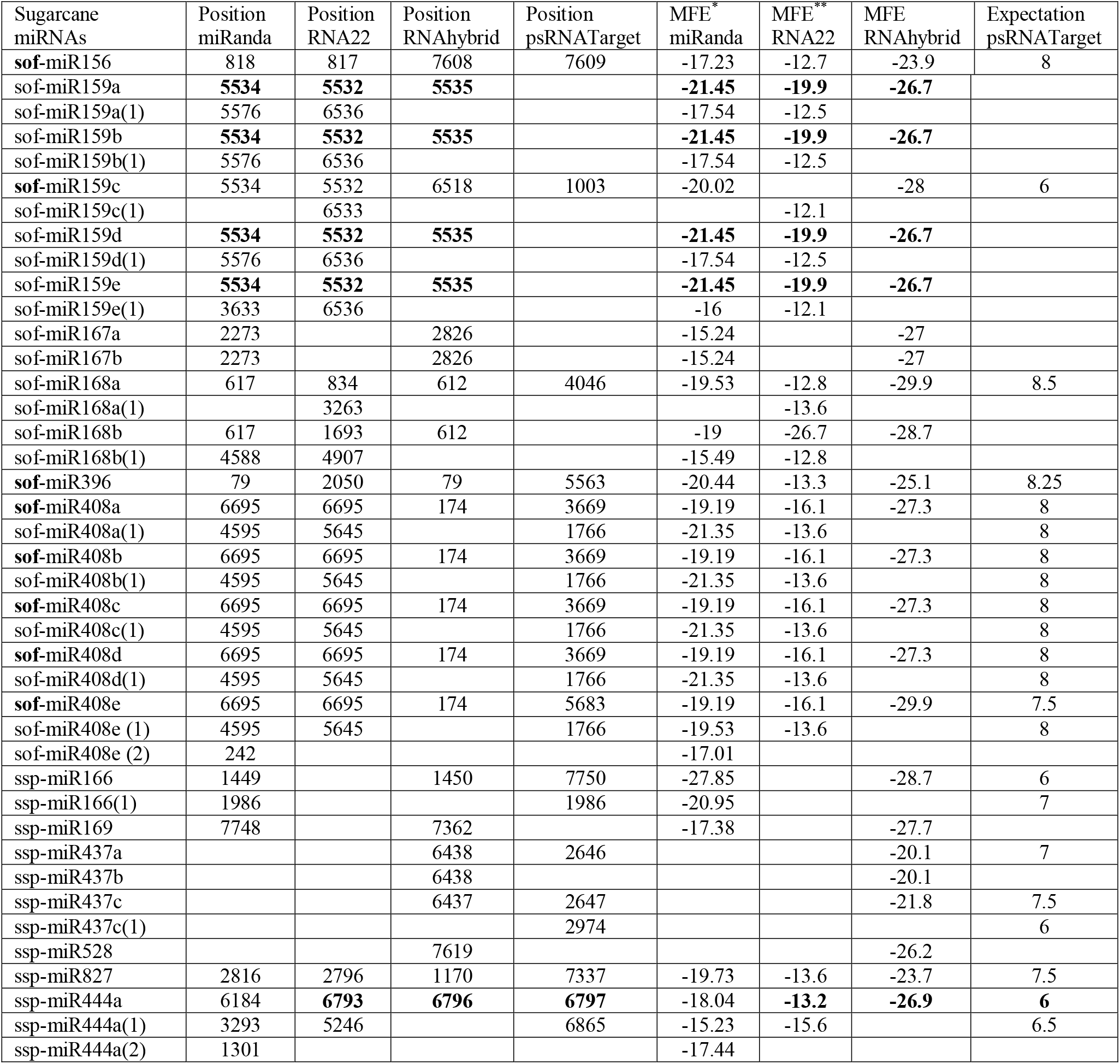
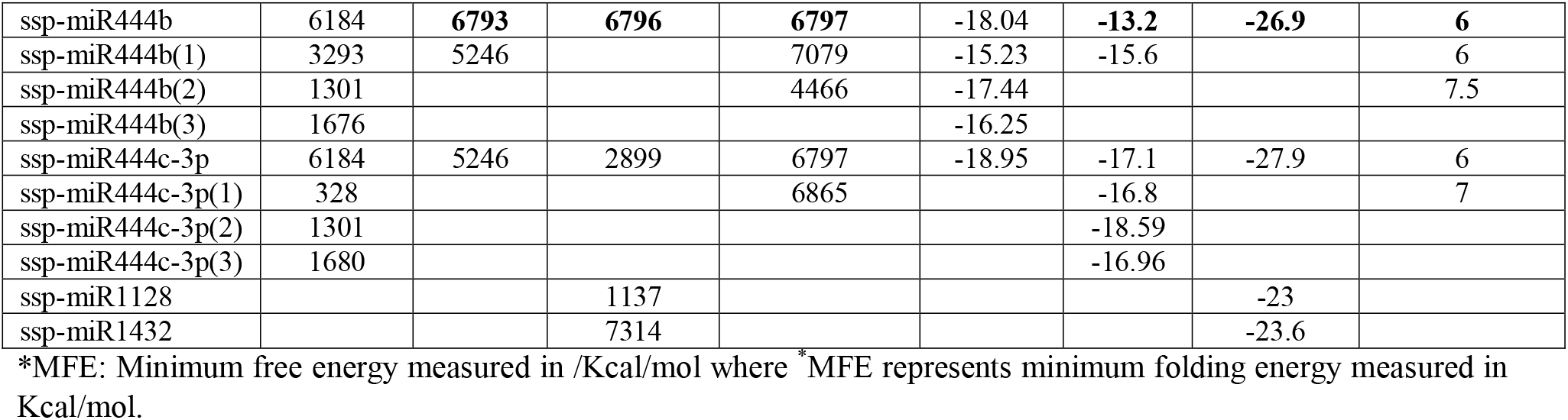
Sugarcane miRNAs and their target positions in the SCBV identified by algorithms.

### 3.7. Predicting Consensus Sugarcane miRNAs for Silencing SCBV Genome

Out of 28 sugarcane miRNAs, only six sugarcane miRNA (sof-miR159 (a, b, d and e) at common locus position 5535 and ssp-miR444 (a, b) at locus 6797) were predicted at the common locus by at least three algorithms used (Fig 7, Table 2 and SF1). Out of 14 consensus miRNAs, only one miRNA of *S.officinarum* (sof-miR159e at locus 5535) with MFE −26.7 Kcal/mol, was considered as the top effective candidate; deliver more efficient silencing of the SCBV genome. The efficacy of sof-miR159e target against SCBV was validated by suppression of RNAi-mediated viral combat through cleavage of viral mRNA or translational inhibition. Multiple loci interactions were observed for sof-miR159e at nucleotide positions 5534-5552 (consensus of three algorithms; miRanda, RNA22 and RNAhybrid) and 2647 (psRNATarget) of ORF3.

**Fig 7.**
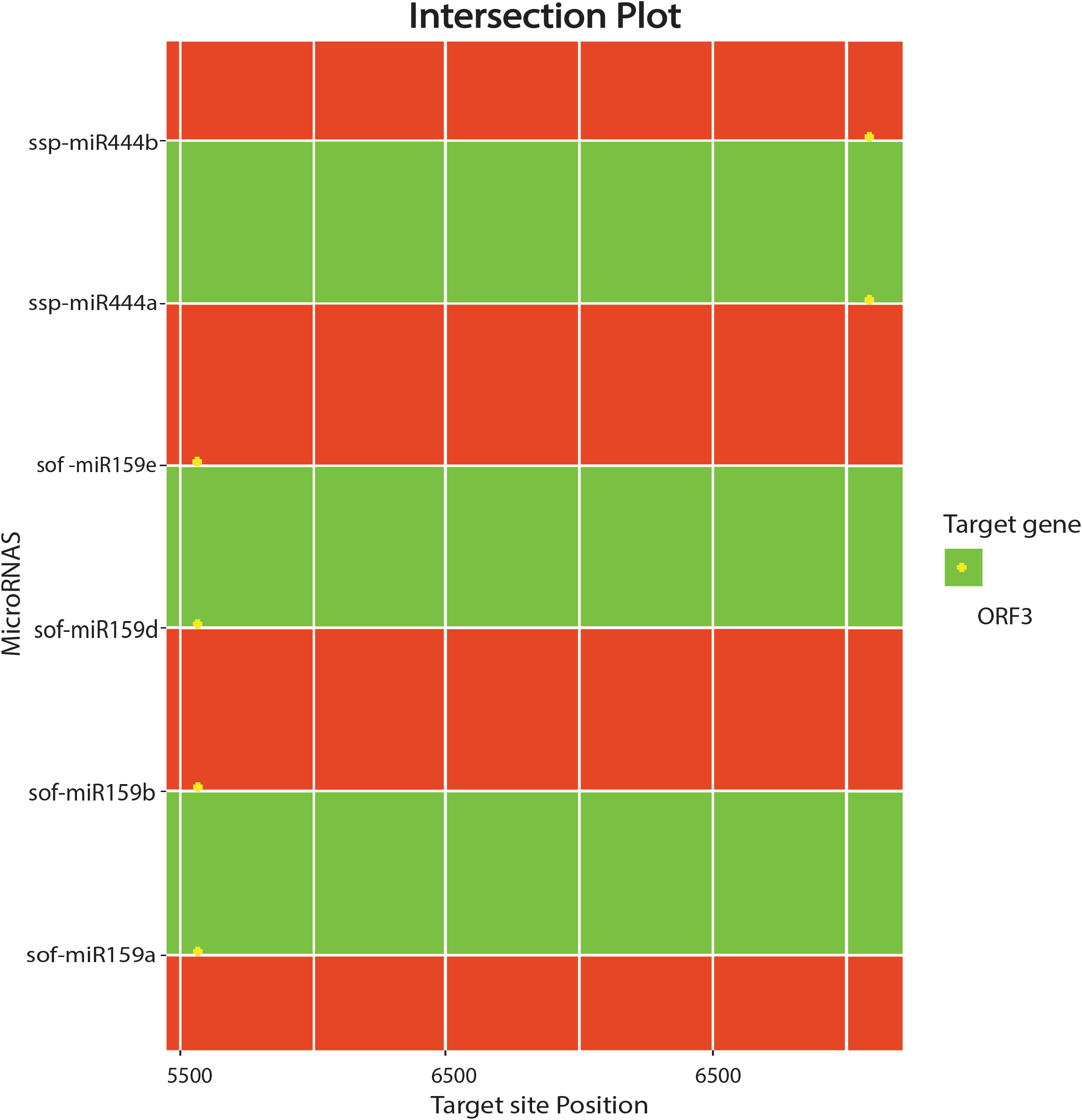
Intersection plot of sugarcane miRNAs predicted from at least three algorithms. Intersection plot was created miRNAs, predicted from at least three algorithms: (miRanda, RNA22 and RNAhybrid). The color code has been given with the figure.

### 3.8. Prediction of Consensus Secondary Structures

The validation of consensus sugarcane-miRNAs was confirmed by the prediction of their stable secondary structures using RNAfold algorithm. Precursors of mature sugarcane miRNAs were manually curated. MFE is the key factor to determine the stable secondary structures of precursors. All the predicted consensus sugarcane miRNA precursors were observed to possess a lower MFE value (ranging from −57.70 to −114.70 kcal/mol) (Table 3).

**Table 3.** Salient parameters of precursor miRNAs was determined and estimation of free energy.

| miRNA ID | Length<br>miRNA | Length<br>precursor | MFE<sup>1</sup><br>(Kcal/mol) | AMFE<sup>2</sup> | MFEI<sup>3</sup> | (G+C)<br>% | $\Delta G$<sup>4</sup><br>(Kcal/mol) |
| --- | --- | --- | --- | --- | --- | --- | --- |
| sof-miR156 | 20 | 137 | -66.20 | -48.32 | -0.96 | 50.00 | -14.30 |
| sof-miR159c | 21 | 238 | -110.60 | -46.47 | -0.88 | 52.38 | -19.70 |
| sof-miR159e | 21 | 264 | -107.50 | -40.71 | -1.06 | 38.09 | -20.40 |
| sof-miR168a | 21 | 104 | -66.20 | -63.65 | -1.02 | 61.90 | -18.20 |
| sof-miR396 | 21 | 134 | -67.40 | -50.29 | -1.17 | 42.85 | -19.60 |
| sof-miR408a | 21 | 283 | -114.70 | -40.53 | -0.60 | 66.66 | -16.00 |
| sof-miR408b | 21 | 286 | -113.20 | -39.58 | -0.59 | 66.66 | -16.00 |
| sof-miR408c | 21 | 286 | -115.80 | -40.48 | -0.60 | 66.66 | -16.00 |
| sof-miR408d | 21 | 215 | -79.00 | -36.76 | -0.55 | 66.66 | -16.00 |
| sof-miR408e | 21 | 283 | -99.00 | -34.98 | -0.56 | 61.90 | -16.00 |
| ssp-miR827 | 21 | 130 | -64.00 | -49.23 | -1.29 | 38.09 | -17.90 |
| ssp-miR444a | 21 | 105 | -57.70 | -54.95 | -1.15 | 47.62 | -14.50 |
| ssp-miR444b | 21 | 106 | -63.70 | -60.09 | -1.26 | 47.62 | -14.50 |
| ssp-miR444c | 21 | 108 | -61.80 | -57.22 | -1.33 | 42.85 | -15.30 |

<sup>1</sup>MFE is minimum free energy. <sup>2</sup>AMFE represents adjusted minimum free energy. <sup>3</sup>MFEI defines as minimum free energy index. <sup>4</sup> $\Delta G$ represents minimum free energy of duplex formation.

The predicted secondary structures of six precursors of pre-miRNAs are shown (Fig 8) as predicted by intersection of three consensus algorithms at the same locus. Top stable secondary structure of sof-MIR159e precursor was predicted with standard features (MFE: 107.50 Kcal/mol, MFEI: 1.06 Kcal/mol). The predicted secondary structures of 14 consensus sugarcane miRNAs passed the standard criteria as mentioned. We have determined the salient characteristics of six consensus precursor miRNAs in this study such as MFE, AMFE, MFEI, length precursor and GC contents. In our studies, length precursor range from 105-266 nucleotides, MFE (−57.70 to −110.70 kcal/mol), AMFE (−39.92 to 60.09) and GC content (38–47%) and MFEI ranges from −0.83 to −1.26.

**Fig 8.**
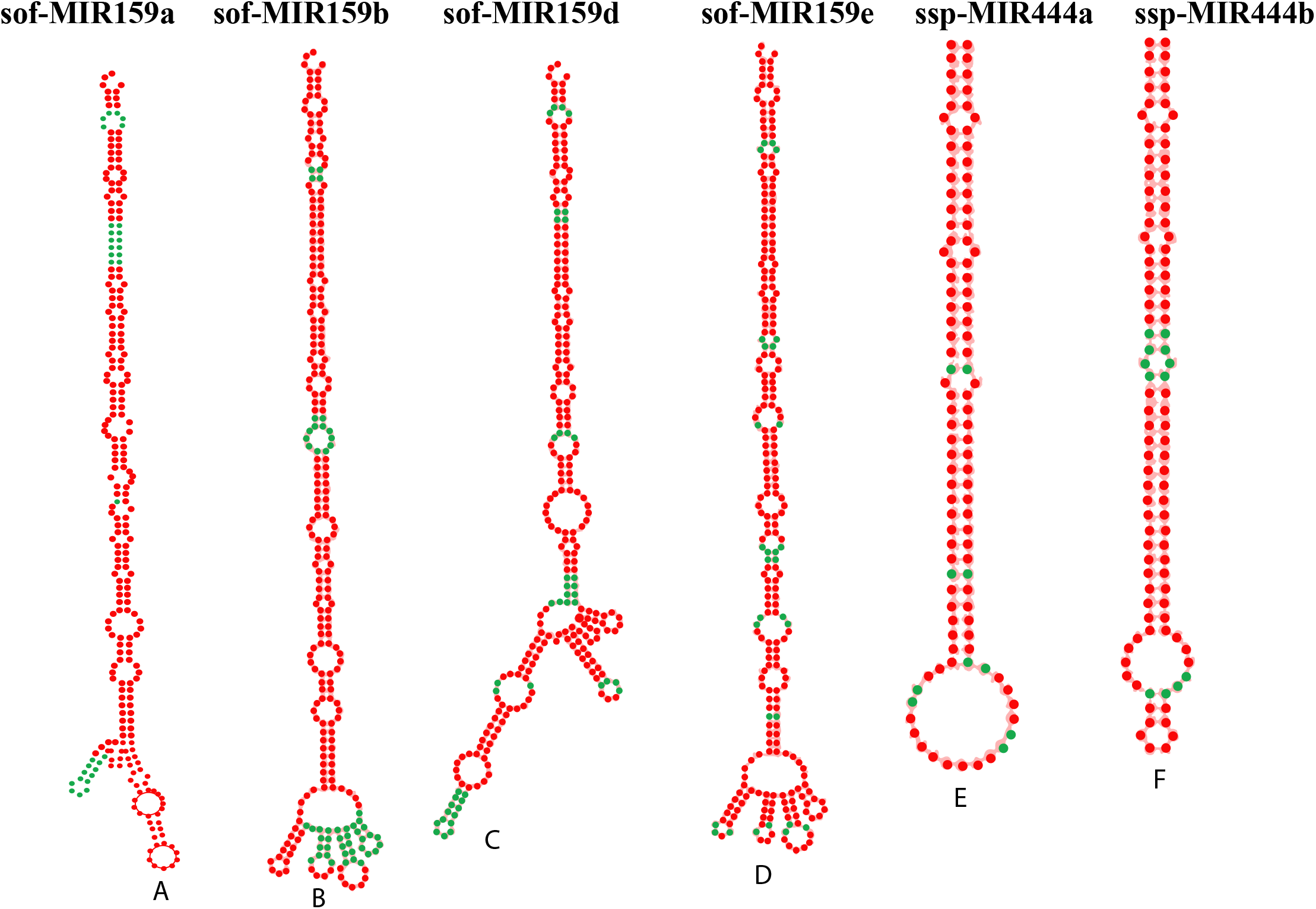
Prediction of secondary structure of stem-loop sequences of sugarcane miRNAs. Six pre-miRNA secondary structures, precursor of sugarcane miRNAs were identified in this study by a consensus of three algorithms. Sugarcane mature miRNA name IDs, accession IDs MFE and MFEI are given as follow: A) sof-MIR159a (MI0001756), −110.30 kcal/mol, −0.87 B) sof-MIR159b (MI0001757), −110.30 kcal/mol, −0.87 C) sof-MIR159d (MI0001758), −105.80 kcal/mol, −0.83 D) sof-MIR159e (MI0001759), −107.50 kcal/mol, −1.06 E) ssp-MIR444a (MI0018185), −57.70 kcal/mol, −1.15 F) ssp-MIR444b (MI0018186), −63.70kcal/mol, −1.26.

### 3.9. Assessment of Free Energy (ΔG) of miRNA-mRNA Interaction

The predicted consensus sugarcane miRNAs were validated by estimating free-energies of miRNA/target duplexes (Table 3). The free-energies (ΔG) of six consensus sugarcane miRNAs are estimated: sof-miR159 (a, b, d) (ΔG: −20.10 kcal/mol), sof-miR15e (ΔG: −20.40 kcal/mol), and ssp-miR444 (a, b) (ΔG: −14.50 kcal/mol).

## 4. Discussion

For Filtering of false-positive results, we study the performance of computational algorithms to validate the miRNA target prediction data. We designed the most effective approach for validation of miRNA target prediction results at individual, union and intersection levels. Computational prediction algorithms offer a rapid method to predict potential host-derived miRNA targets in the virus genome. We evaluated the efficiency of four frequently used computational tools cited in literature—miRanda, RNA22, RNAhybrid and psRNATarget to identify miRNA-target prediction and to analyze miRNA-mRNA interactions as summarized in (Table1). miRanda is widely used algorithms, includes main aspects of miRNA-target prediction; conservation level and miRNA 3’ site [46]. RNA22 adopts a sequence-based pattern recognition mechanism to identify potential targets independently of conservation status and also include 3’ UTR, 5’UTR and CDS regions for target prediction [34]. RNA22 algorithm is a novel alternative option for exploring new miRNA: mRNA interactions, because of its unique capabilities but likelihood of high rate of potential false-positive [47]. RNAhybrid is widely used to find the energetically most favorable hybridization binding sites [36, 48]. We calculate the MFE and determine the target inhibition as recommended from Broderson conclusion using RNAhybrid [37]. psRNATarget is a novel plant candidate miRNAs prediction algorithm used for assessment of miRNA-mRNA interaction [38].

Several potential sugarcane miRNA targets and miRNA-mRNA interaction could be consensually predicted by all the algorithms (Fig 7). miRanda and psRNATarget are two powerful plant miRNA prediction algorithms for identifying hybridization sites in the viral genome. Plant miRNAs are responsible to induce the degradation of the target gens using perfect or imperfect complementarity base pairing [49]. The current study demonstrates that SCBV genome components (ORF1, ORF2 and ORF3) are susceptible to be targeted by a set of consensus sugarcane miRNAs. In addition, sof-miR159 (a, b, d and e) was found to target the ORF3 at consensus hybridization site by at least three algorithms (Fig 8). Free energy assessment is a dynamic feature of miRNA and target binding. Previous study revealed a significant correlation of free energy between the translational repression and the hybridization binding of the seed region [50]. The thermodynamic stability of the miRNA-mRNA duplex was estimated by assessment of free energy to monitor site accessibility for determination of the secondary structure duplex [27]. In order to validate miRNA-mRNA interaction, free energy of duplex was assessed as shown in (Table 2). Our prediction results are outstanding to show high stability of sugarcane-encoded miRNA-SCBV-mRNA duplex at low level of free energy at low free energy level as shown in the (Table 3 and Fig 8). RNA duplex is considered more stable due to stronger binding of miRNA to mRNA [51, 52].

We used union and intersection approaches to reduce false-positive prediction. Union approach relies on combining more than one target prediction tool: true and false targets. Sensitivity level of predicted targets increases due to decrease of specificity. Intersection approach is entirely different and depends upon the combining of two or more computational algorithmic tools which enhances the specificity level of predicted targets due to decrease of sensitivity [53]. Our target prediction results revealed that both the computational approaches achieved the best outcome with maximum of performance for predicting and estimating the best targets as shown in (Fig. 6 and 7). Previous studies have also reported the silencing of plant viruses using host-derived miRNAs applying a set of computational algorithms. Identification and evaluation of best-fit candidate miRNA targets of different crops plants was concluded successfully in PVY [54], maize chlorotic mottle virus (MCMV) [55], CLCuKoV-Bu [56], RYMV [57] and SCBGAV to find miRNA-target interaction [58]. We have designed the same novel bioinformatics approach for target prediction in the SCBV genome to control the emerging badnavirus in sugarcane cultivars.

In our previous study, we identified the most potential consensus sugarcane miRNA (sof-miR396) to target ORF3 of SCBGAV genome using multiple computational algorithms [58]. The quantity of false-positive miRNA-target interaction estimated by multiple algorithms depends upon the mode of miRNA-target recognition. MFE is also another one of the important factors to affect the miRNA-target interaction for result validation [59]. To set a lower MFE value, give rise high probability of miRNA-target complex formation [60]. In the current study, for miRanda analysis, a stringent cut-off −15 kcal/mol was set for narrowing down the miRNA candidates. Similarly to validate host-virus interaction, a MFE cut-off −20 kcal/mol applied for RNAhybrid analysis [32].

In the previous study, consensus prediction results were sorted based on a stringent cut-off −20 kcal/mol by intersection of two potential algorithms to explore possible interaction between *Aedes aegypti* cellular miRNA and arboviruses [61]. Although MFE has considerable role for development miRNA-mRNA complex but it does not certify that the interaction will lead to functional changes. In the current study, we identify six potential miRNA hybridization binding sites that have exhibited low MFE and free-energy of duplex formation. These predicted miRNAs not only have potential target SCBV genome at transgenic level but also have stronger probability to develop miRNA-viral mRNA complex formation. These miRNAs also have chance to participate in SCBV replication mechanism: a consensus sugarcane miRNA (sof-miR396) has binding site within the SCBV large intergenic region (LIR) at locus 79 predicted by miRanda and RNAhybrid algorithms. In the previous study, we predicted that the sof-miRNA396 is an effective candidate to target SCBGAV genome [58]. The sof-miR159e was predicted by all the algorithms. The miR159 was explored to have strong role for silencing of *GAMYB* to enable normal growth [62]. Phe-MIR159 was involved to regulate gene responsible to secondary thickening in *Phyllostachys edulis* plant [63]. It is important to assess the function of predicted potential consensus miRNAs for identification of badnavirus replication to demonstrate SCBV replication experimentally. A hypothetical model was designed to show sugarcane-derived miRNAs can inhibit SCBV mRNA and sugarcane genes against SCBV virus as shown in (Fig 9). It facilitates to plant-encoded miRNAs to cleavage of SCBV miRNA.

**Fig 9.**
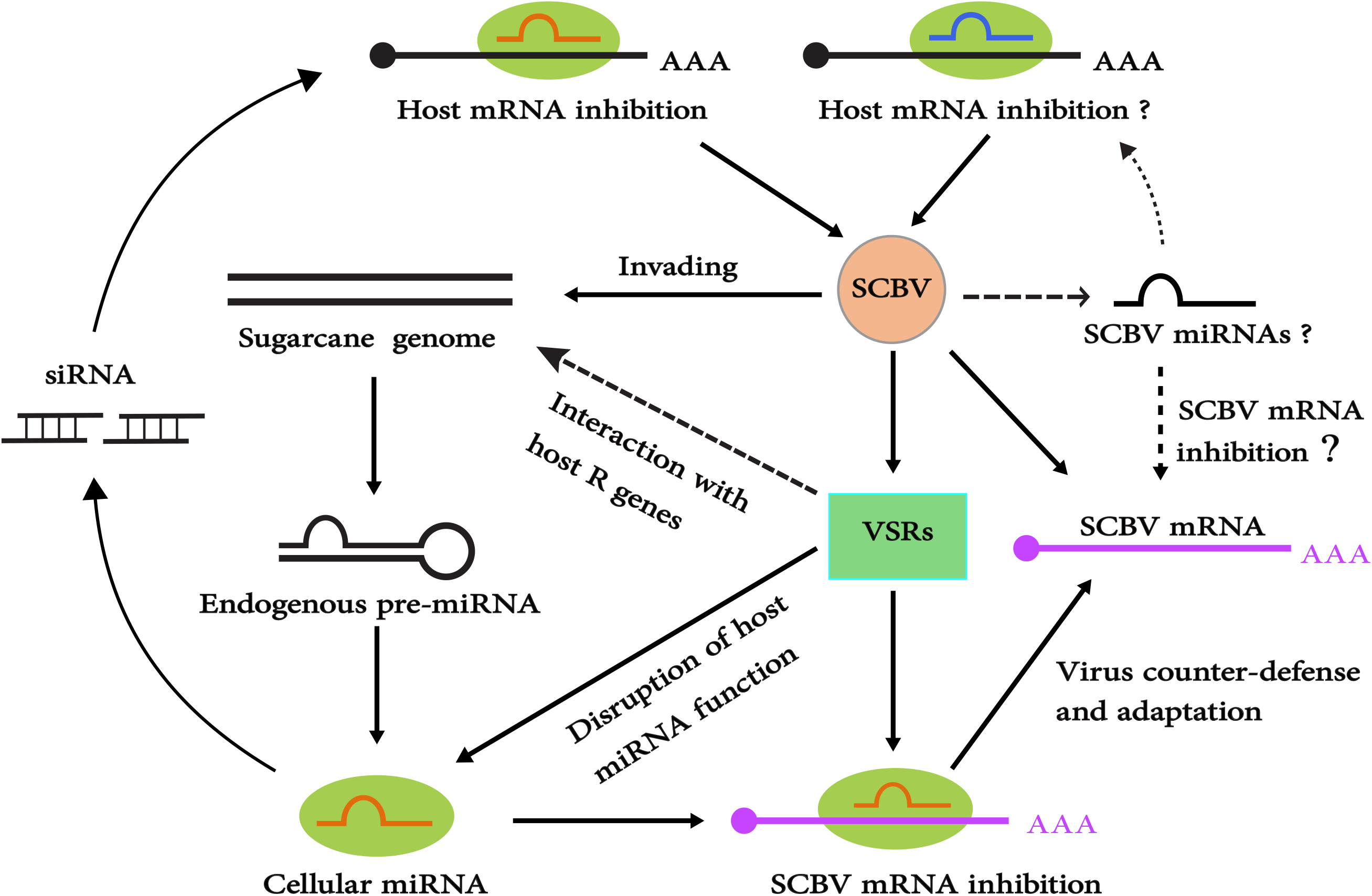
Schematic model designed for amiRNA-based silencing mechanism. amiRNA based gene silencing mechanism was designed in schematic manner. SCBV can activate the production of sugarcane endogenous miRNAs after infection. The sugarcane miRNAs can target SCBV mRNA for degradation.

RNAi screens is novel technology to discover various cellular functions and to identify host-derived factors of viruses [64]. We selected 28 experimental validated sugarcane miRNAs which had annotated targets that are part of the SCBV. We have designed amiRNA-based gene construct of SCBV which contains modified miRNA/miRNA* sequence in duplex of precursor (sof-MIR159) with artificial sequence to create mature amiRNA as shown in (Fig 10). The amiRNA-based silencing technology is successfully validated in many crop plants to control emerging plant viruses [18, 19, 21]. In summary, our computational work for the silencing of the SCBV genome could offer new approach for the remodel of existing strategy as antiviral agents. We demonstrated to minimize novel antiviral effects of host-derived miRNAs against SCBV.

**Fig 10.**
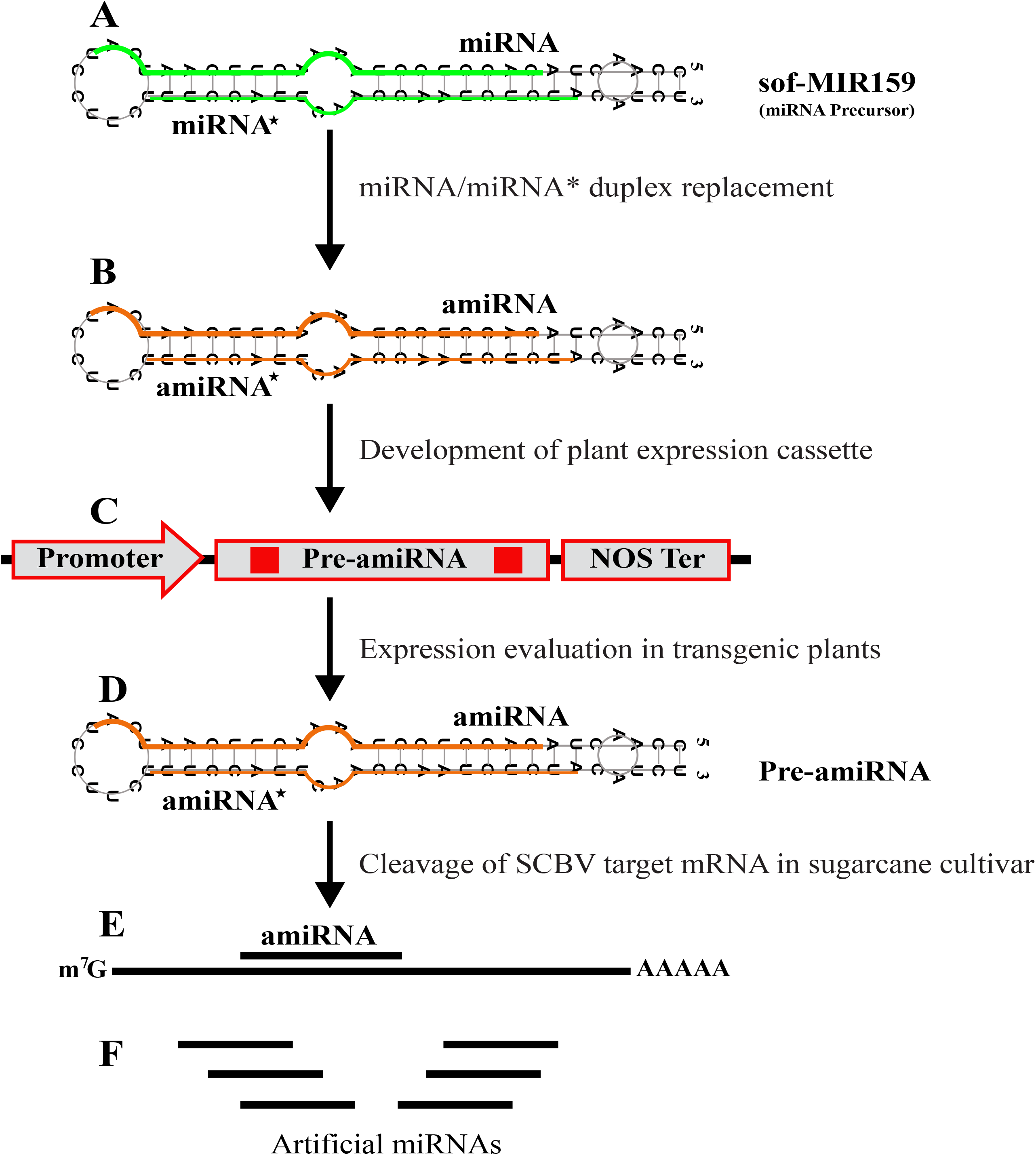
Systematic mechanism of amiRNA-mediated gene silencing in transgenic sugarcane. Systematic mechanism was designed to impart amiRNA-mediated gene silencing in transgenic sugarcane plants. Candidate amiRNA is implanted into sof-MIR159 precursor after miRNA/miRNA duplex replacement. This pre-amiRNA is processed to generate mature amiRNA/amiRNA* duplex.

## 5. Conclusions

The SCBV has appeared as a major problem in China. SCBV diminishes the quantitative yield in all sugarcane cultivars. In the current study, we applied computational tools to predict and comprehensively analyse the candidate miRNA from sugarcane against the SCBV, prior to cloning. Among them, sof-miR159e was predicted as top effective candidate that could target the vital gene (ORF3) of the SCBV genome. Our results conclude an alternative strategy to existing molecular approaches could be re-purposed to control badnaviral infections. The current findings provide a proof of novel scheme to construct amiRNA-based genome silencing therapeutics to combat SCBV.

## Supporting information

File S1: Computational prediction of Sugarcane-encoded microRNA targets against SCBV genome using four algorithms.

Table S1: Salient features of sugarcane microRNAs retrieved from the miRBase database.

Table S2: Prediction of sugarcane microRNAs in the SCBV genome.

Table S3: Sugarcane-encoded miRNA target binding sites locus position in each gene of the SCBV genome.

## Supplementary Materials

The following are available (Table S1, S2, S3 and File S1).

## Acknowledgments

We are highly thankful to our lab colleagues for their assistance in data analysis. We wish to thank Dr. Zhiqiang Xia (ITBB) for providing facilities and assistance in the construction of the Circos plot.

## Author Contributions

Conceptualization, M.A.A. and S.Z.; Methodology, M.A.A., X.H and F.A; Software, M.A.A. F.A and X.H; Validation, F.A, L.S and X.F.; Formal Analysis, M.A.A, X.H.; Investigation, M.A.A., L.S and X.F.; Resources, S.Z. and X.F.; Data Curation, S.Z, L.S. and X.F; Writing—Original Draft Preparation, M.A.A and F.A; Writing—Review and Editing, M.A.A, F.A and S.Z.; Visualization, L.S; Supervision, S.Z.; Funding Acquisition, S.Z, X.F and X.H. All authors have read and agree to the published version of the manuscript.

