## Supplementary material for "An Algorithmic framework for genome-wide identification of Sugarcane (*Saccharum officinarum* L.)-encoded microRNA targets against SCBV": File S1: Computational prediction of Sugarcane-encoded microRNA targets against SCBV genome using four algorithms.

```
=-=-=-=-=-=-=-=-=-=-=-=-=-=-=-=-=-=-=-=-=-=-=-=-=-=-=-=-=-=-=-=-=
miranda v3.3a    microRNA Target Scanning Algorithm
=-=-=-=-=-=-=-=-=-=-=-=-=-=-=-=-=-=-=-=-=-=-=-=-=-=-=-=-=-=-=-=-=
(c) 2003 Memorial Sloan-Kettering Cancer Center, New York

Authors: Anton Enright, Bino John, Chris Sander and Debora Marks
(mirnatargets (at) cbio.mskcc.org - reaches all authors)

Software written by: Anton Enright
Distributed for anyone to use under the GNU Public License (GPL),
See the files 'COPYING' and 'LICENSE' for details

If you use this software please cite:
Enright AJ, John B, Gaul U, Tuschl T, Sander C and Marks DS;
(2003) Genome Biology; 5(1):R1.

   miranda comes with ABSOLUTELY NO WARRANTY;
   This is free software, and you are welcome to redistribute it
   under certain conditions; type `miranda --license' for details.

Current Settings:
=-=-=-=-=-=-=-=-=-=-=-=-=-=-=-=-=-=-=-=-=-=-=-=-=-=-=-=-=-=-=-=-=
Query Filename:   ./seq/mirna/sof_mirna.fasta
Reference Filename:     ./seq/gene/sof/JN377537.fasta
Gap Open Penalty: -9.000000
Gap Extend Penalty:     -4.000000
Score Threshold:  130.000000
Energy Threshold: -15.000000 kcal/mol
Scaling Parameter:      4.000000
=-=-=-=-=-=-=-=-=-=-=-=-=-=-=-=-=-=-=-=-=-=-=-=-=-=-=-=-=-=-=-=-=
Read Sequence:sof-miR156 (20 nt)
Read Sequence:JN377537.1 Sugarcane bacilliform virus isolate BRU, complete
genome(7884 nt)
=-=-=-=-=-=-=-=-=-=-=-=-=-=-=-=-=-=-=-=-=-=-=-=-=-=-=-=-=-=-=-=-=
Performing Scan: sof-miR156 vs JN377537.1
=-=-=-=-=-=-=-=-=-=-=-=-=-=-=-=-=-=-=-=-=-=-=-=-=-=-=-=-=-=-=-=-=

   Forward: Score: 150.000000  Q:2 to 15  R:818 to 837 Align Len (13) (69.23%)
(92.31%)

   Query:    3' cacgagUGAGAGAAGACAGu 5'
                   ::| |:|||||||
   Ref:      5' agcagaGTTGTTTTCTGTCa 3'

   Energy:  -17.230000 kCal/Mol

Scores for this hit:
>sof-miR156 JN377537.1  150.00      -17.23      2 15  818 837     13    69.23%
      92.31%

Score for this Scan:
Seq1,Seq2,Tot Score,Tot Energy,Max Score,Max Energy,Strand,Len1,Len2,Positions
>>sof-miR156      JN377537.1  150.00      -17.23      150.00      -17.23      1
      20    7884   818
Complete

Read Sequence:sof-miR159a (21 nt)
Read Sequence:JN377537.1 Sugarcane bacilliform virus isolate BRU, complete
genome(7884 nt)
=-=-=-=-=-=-=-=-=-=-=-=-=-=-=-=-=-=-=-=-=-=-=-=-=-=-=-=-=-=-=-=-=
Performing Scan: sof-miR159a vs JN377537.1
=-=-=-=-=-=-=-=-=-=-=-=-=-=-=-=-=-=-=-=-=-=-=-=-=-=-=-=-=-=-=-=-=
```

```
   Forward: Score: 144.000000  Q:2 to 20  R:5534 to 5552 Align Len (18) (72.22%)
(83.33%)

   Query:    3' guCUCGAGGGAAGUUAGGUUu 5'
                 |||||  : ||||||||:
   Ref:      5' ttGAGCT--TATCAATCCAGa 3'

   Energy:  -21.450001 kCal/Mol

Scores for this hit:
>sof-miR159a      JN377537.1  144.00      -21.45      2 20  5534 5552   18
      72.22%      83.33%


   Forward: Score: 139.000000  Q:2 to 16  R:5576 to 5596 Align Len (14) (71.43%)
(85.71%)

   Query:    3' gucucgAGGGAAGUUAGGUUu 5'
                     | |:| |||||||:
   Ref:      5' ttacaaTGCTTACAATCCAGa 3'

   Energy:  -17.549999 kCal/Mol

Scores for this hit:
>sof-miR159a      JN377537.1  139.00      -17.55      2 16  5576 5596   14
      71.43%      85.71%

Score for this Scan:
Seq1,Seq2,Tot Score,Tot Energy,Max Score,Max Energy,Strand,Len1,Len2,Positions
>>sof-miR159a     JN377537.1  283.00      -39.00      144.00      -21.45      2
      21    7884   5534 5576
Complete

Read Sequence:sof-miR159b (21 nt)
Read Sequence:JN377537.1 Sugarcane bacilliform virus isolate BRU, complete
genome(7884 nt)
=-=-=-=-=-=-=-=-=-=-=-=-=-=-=-=-=-=-=-=-=-=-=-=-=-=-=-=-=-=-=-=-=
Performing Scan: sof-miR159b vs JN377537.1
=-=-=-=-=-=-=-=-=-=-=-=-=-=-=-=-=-=-=-=-=-=-=-=-=-=-=-=-=-=-=-=-=

   Forward: Score: 144.000000  Q:2 to 20  R:5534 to 5552 Align Len (18) (72.22%)
(83.33%)

   Query:    3' guCUCGAGGGAAGUUAGGUUu 5'
                 |||||  : ||||||||:
   Ref:      5' ttGAGCT--TATCAATCCAGa 3'

   Energy:  -21.450001 kCal/Mol

Scores for this hit:
>sof-miR159b      JN377537.1  144.00      -21.45      2 20  5534 5552   18
      72.22%      83.33%


   Forward: Score: 139.000000  Q:2 to 16  R:5576 to 5596 Align Len (14) (71.43%)
(85.71%)

   Query:    3' gucucgAGGGAAGUUAGGUUu 5'
                     | |:| |||||||:
   Ref:      5' ttacaaTGCTTACAATCCAGa 3'

   Energy:  -17.549999 kCal/Mol
```

```
Scores for this hit:
>sof-miR159b      JN377537.1  139.00      -17.55      2 16  5576 5596   14
      71.43%      85.71%

Score for this Scan:
Seq1,Seq2,Tot Score,Tot Energy,Max Score,Max Energy,Strand,Len1,Len2,Positions
>>sof-miR159b     JN377537.1  283.00      -39.00      144.00      -21.45      3
      21    7884   5534 5576
Complete

Read Sequence:sof-miR159c (21 nt)
Read Sequence:JN377537.1 Sugarcane bacilliform virus isolate BRU, complete
genome(7884 nt)
=-=-=-=-=-=-=-=-=-=-=-=-=-=-=-=-=-=-=-=-=-=-=-=-=-=-=-=-=-=-=-=
Performing Scan: sof-miR159c vs JN377537.1
=-=-=-=-=-=-=-=-=-=-=-=-=-=-=-=-=-=-=-=-=-=-=-=-=-=-=-=-=-=-=-=

   Forward: Score: 144.000000  Q:2 to 20  R:5534 to 5552 Align Len (18) (72.22%)
(83.33%)

   Query:    3' ucCUCGAGGGAAGUUAGGUUc 5'
                  |||||  : ||||||||:
   Ref:      5' ttGAGCT--TATCAATCCAGa 3'

   Energy:  -20.020000 kCal/Mol

Scores for this hit:
>sof-miR159c      JN377537.1  144.00      -20.02      2 20  5534 5552   18
      72.22%      83.33%

Score for this Scan:
Seq1,Seq2,Tot Score,Tot Energy,Max Score,Max Energy,Strand,Len1,Len2,Positions
>>sof-miR159c     JN377537.1  144.00      -20.02      144.00      -20.02      4
      21    7884   5534
Complete

Read Sequence:sof-miR159d (21 nt)
Read Sequence:JN377537.1 Sugarcane bacilliform virus isolate BRU, complete
genome(7884 nt)
=-=-=-=-=-=-=-=-=-=-=-=-=-=-=-=-=-=-=-=-=-=-=-=-=-=-=-=-=-=-=-=
Performing Scan: sof-miR159d vs JN377537.1
=-=-=-=-=-=-=-=-=-=-=-=-=-=-=-=-=-=-=-=-=-=-=-=-=-=-=-=-=-=-=-=

   Forward: Score: 144.000000  Q:2 to 20  R:5534 to 5552 Align Len (18) (72.22%)
(83.33%)

   Query:    3' guCUCGAGGGAAGUUAGGUUu 5'
                  |||||  : ||||||||:
   Ref:      5' ttGAGCT--TATCAATCCAGa 3'

   Energy:  -21.450001 kCal/Mol

Scores for this hit:
>sof-miR159d      JN377537.1  144.00      -21.45      2 20  5534 5552   18
      72.22%      83.33%


   Forward: Score: 139.000000  Q:2 to 16  R:5576 to 5596 Align Len (14) (71.43%)
(85.71%)

   Query:    3' gucucgAGGGAAGUUAGGUUu 5'
                      | |:| |||||||:
   Ref:      5' ttacaaTGCTTACAATCCAGa 3'
```

```
   Energy:  -17.549999 kCal/Mol

Scores for this hit:
>sof-miR159d      JN377537.1  139.00      -17.55      2 16  5576 5596   14
      71.43%      85.71%

Score for this Scan:
Seq1,Seq2,Tot Score,Tot Energy,Max Score,Max Energy,Strand,Len1,Len2,Positions
>>sof-miR159d     JN377537.1  283.00      -39.00      144.00      -21.45      5
      21    7884   5534 5576
Complete

Read Sequence:sof-miR159e (21 nt)
Read Sequence:JN377537.1 Sugarcane bacilliform virus isolate BRU, complete
genome(7884 nt)
=-=-=-=-=-=-=-=-=-=-=-=-=-=-=-=-=-=-=-=-=-=-=-=-=-=-=-=-=-=-=-=-=
Performing Scan: sof-miR159e vs JN377537.1
=-=-=-=-=-=-=-=-=-=-=-=-=-=-=-=-=-=-=-=-=-=-=-=-=-=-=-=-=-=-=-=-=

   Forward: Score: 148.000000  Q:2 to 20  R:5534 to 5552 Align Len (18) (77.78%)
(83.33%)

   Query:    3' uuCUCGAGGAAAGUUAGGUUu 5'
                  |||||  | ||||||||:
   Ref:      5' ttGAGCT--TATCAATCCAGa 3'

   Energy:  -21.450001 kCal/Mol

Scores for this hit:
>sof-miR159e      JN377537.1  148.00      -21.45      2 20  5534 5552   18
      77.78%      83.33%


   Forward: Score: 134.000000  Q:2 to 17  R:3633 to 3656 Align Len (18) (66.67%)
(83.33%)

   Query:    3' uucucGAGG--AA-AGUUAGGUUu 5'
                     :|||  || |:||||||:
   Ref:      5' aggctTTCCGGTTGTTAATCCAGt 3'

   Energy:  -16.000000 kCal/Mol

Scores for this hit:
>sof-miR159e      JN377537.1  134.00      -16.00      2 17  3633 3656   18
      66.67%      83.33%

Score for this Scan:
Seq1,Seq2,Tot Score,Tot Energy,Max Score,Max Energy,Strand,Len1,Len2,Positions
>>sof-miR159e     JN377537.1  282.00      -37.45      148.00      -21.45      6
      21    7884   5534 3633
Complete

Read Sequence:sof-miR167a (21 nt)
Read Sequence:JN377537.1 Sugarcane bacilliform virus isolate BRU, complete
genome(7884 nt)
=-=-=-=-=-=-=-=-=-=-=-=-=-=-=-=-=-=-=-=-=-=-=-=-=-=-=-=-=-=-=-=-=
Performing Scan: sof-miR167a vs JN377537.1
=-=-=-=-=-=-=-=-=-=-=-=-=-=-=-=-=-=-=-=-=-=-=-=-=-=-=-=-=-=-=-=-=

   Forward: Score: 137.000000  Q:3 to 20  R:2273 to 2292 Align Len (17) (70.59%)
(82.35%)

   Query:    3' guCUAGUACGACCGUCGAAgu 5'
                  ||| :||:|  ||||||
```

```
   Ref:      5' ttGATGGTGTT-ACAGCTTat 3'

   Energy:  -15.240000 kCal/Mol

Scores for this hit:
>sof-miR167a      JN377537.1  137.00      -15.24      3 20  2273 2292   17
      70.59%      82.35%

Score for this Scan:
Seq1,Seq2,Tot Score,Tot Energy,Max Score,Max Energy,Strand,Len1,Len2,Positions
>>sof-miR167a     JN377537.1  137.00      -15.24      137.00      -15.24      7
      21    7884   2273
Complete

Read Sequence:sof-miR167b (21 nt)
Read Sequence:JN377537.1 Sugarcane bacilliform virus isolate BRU, complete
genome(7884 nt)
=-=-=-=-=-=-=-=-=-=-=-=-=-=-=-=-=-=-=-=-=-=-=-=-=-=-=-=-=-=-=-=-=
Performing Scan: sof-miR167b vs JN377537.1
=-=-=-=-=-=-=-=-=-=-=-=-=-=-=-=-=-=-=-=-=-=-=-=-=-=-=-=-=-=-=-=-=

   Forward: Score: 137.000000  Q:3 to 20  R:2273 to 2292 Align Len (17) (70.59%)
(82.35%)

   Query:    3' guCUAGUACGACCGUCGAAgu 5'
                  ||| :||:|  ||||||
   Ref:      5' ttGATGGTGTT-ACAGCTTat 3'

   Energy:  -15.240000 kCal/Mol

Scores for this hit:
>sof-miR167b      JN377537.1  137.00      -15.24      3 20  2273 2292   17
      70.59%      82.35%

Score for this Scan:
Seq1,Seq2,Tot Score,Tot Energy,Max Score,Max Energy,Strand,Len1,Len2,Positions
>>sof-miR167b     JN377537.1  137.00      -15.24      137.00      -15.24      8
      21    7884   2273
Complete

Read Sequence:sof-miR168a (21 nt)
Read Sequence:JN377537.1 Sugarcane bacilliform virus isolate BRU, complete
genome(7884 nt)
=-=-=-=-=-=-=-=-=-=-=-=-=-=-=-=-=-=-=-=-=-=-=-=-=-=-=-=-=-=-=-=-=
Performing Scan: sof-miR168a vs JN377537.1
=-=-=-=-=-=-=-=-=-=-=-=-=-=-=-=-=-=-=-=-=-=-=-=-=-=-=-=-=-=-=-=-=

   Forward: Score: 142.000000  Q:2 to 16  R:617 to 638 Align Len (15) (80.00%)
(86.67%)

   Query:    3' cagggcUAGACGU-GGUUCGCu 5'
                      ||||| | |||||:|
   Ref:      5' catgaaATCTGAAGCCAAGTGg 3'

   Energy:  -19.530001 kCal/Mol

Scores for this hit:
>sof-miR168a      JN377537.1  142.00      -19.53      2 16  617 638     15
      80.00%      86.67%

Score for this Scan:
Seq1,Seq2,Tot Score,Tot Energy,Max Score,Max Energy,Strand,Len1,Len2,Positions
>>sof-miR168a     JN377537.1  142.00      -19.53      142.00      -19.53      9
      21    7884   617
```

```
Complete

Read Sequence:sof-miR168b (20 nt)
Read Sequence:JN377537.1 Sugarcane bacilliform virus isolate BRU, complete
genome(7884 nt)
=-=-=-=-=-=-=-=-=-=-=-=-=-=-=-=-=-=-=-=-=-=-=-=-=-=-=-=-=-=-=-=-=
Performing Scan: sof-miR168b vs JN377537.1
=-=-=-=-=-=-=-=-=-=-=-=-=-=-=-=-=-=-=-=-=-=-=-=-=-=-=-=-=-=-=-=-=

   Forward: Score: 133.000000  Q:2 to 15  R:617 to 638 Align Len (15) (73.33%)
(80.00%)

   Query:    3' cagggcUAGAC--GGGUUCGCu 5'
                      |||||   |||||:|
   Ref:      5' catgaaATCTGAAGCCAAGTGg 3'

   Energy:  -19.000000 kCal/Mol

Scores for this hit:
>sof-miR168b      JN377537.1  133.00      -19.00      2 15  617 638     15
      73.33%      80.00%


   Forward: Score: 130.000000  Q:3 to 19  R:4588 to 4607 Align Len (16) (62.50%)
(75.00%)

   Query:    3' caGGGCUAGACGGGUUCGcu 5'
                  |::|| |   ||||||
   Ref:      5' gcCTTGAACAAACCAAGCag 3'

   Energy:  -15.490000 kCal/Mol

Scores for this hit:
>sof-miR168b      JN377537.1  130.00      -15.49      3 19  4588 4607   16
      62.50%      75.00%

Score for this Scan:
Seq1,Seq2,Tot Score,Tot Energy,Max Score,Max Energy,Strand,Len1,Len2,Positions
>>sof-miR168b     JN377537.1  263.00      -34.49      133.00      -19.00      10
      20    7884   617 4588
Complete

Read Sequence:sof-miR396 (21 nt)
Read Sequence:JN377537.1 Sugarcane bacilliform virus isolate BRU, complete
genome(7884 nt)
=-=-=-=-=-=-=-=-=-=-=-=-=-=-=-=-=-=-=-=-=-=-=-=-=-=-=-=-=-=-=-=-=
Performing Scan: sof-miR396 vs JN377537.1
=-=-=-=-=-=-=-=-=-=-=-=-=-=-=-=-=-=-=-=-=-=-=-=-=-=-=-=-=-=-=-=-=

   Forward: Score: 130.000000  Q:2 to 20  R:79 to 104 Align Len (23) (65.22%)
(73.91%)

   Query:    3' guCAAGUUCUU-----UCGACACCUu 5'
                  |||:|||:|     ||||||| |
   Ref:      5' gaGTTTAAGGACAACTAGCTGTGCAa 3'

   Energy:  -20.440001 kCal/Mol

Scores for this hit:
>sof-miR396 JN377537.1  130.00      -20.44      2 20  79 104      23    65.22%
      73.91%

Score for this Scan:
Seq1,Seq2,Tot Score,Tot Energy,Max Score,Max Energy,Strand,Len1,Len2,Positions
```

```
>>sof-miR396      JN377537.1  130.00      -20.44      130.00      -20.44      11
      21    7884   79
Complete

Read Sequence:sof-miR408a (21 nt)
Read Sequence:JN377537.1 Sugarcane bacilliform virus isolate BRU, complete
genome(7884 nt)
=-=-=-=-=-=-=-=-=-=-=-=-=-=-=-=-=-=-=-=-=-=-=-=-=-=-=-=-=-=-=-=-=
Performing Scan: sof-miR408a vs JN377537.1
=-=-=-=-=-=-=-=-=-=-=-=-=-=-=-=-=-=-=-=-=-=-=-=-=-=-=-=-=-=-=-=-=

   Forward: Score: 154.000000  Q:2 to 15  R:4595 to 4615 Align Len (13) (84.62%)
(84.62%)

   Query:    3' cggucccUUCUCCGUCACGUc 5'
                       |||  ||||||||
   Ref:      5' acaaaccAAGCAGCAGTGCAg 3'

   Energy:  -21.350000 kCal/Mol

Scores for this hit:
>sof-miR408a      JN377537.1  154.00      -21.35      2 15  4595 4615   13
      84.62%      84.62%


   Forward: Score: 135.000000  Q:2 to 16  R:6695 to 6715 Align Len (14) (71.43%)
(78.57%)

   Query:    3' cggucccUUCUCCGUCACGUc 5'
                      |||  | :||||||
   Ref:      5' ctgtcaGAACTGATAGTGCAg 3'

   Energy:  -19.190001 kCal/Mol

Scores for this hit:
>sof-miR408a      JN377537.1  135.00      -19.19      2 16  6695 6715   14
      71.43%      78.57%

Score for this Scan:
Seq1,Seq2,Tot Score,Tot Energy,Max Score,Max Energy,Strand,Len1,Len2,Positions
>>sof-miR408a     JN377537.1  289.00      -40.54      154.00      -21.35      12
      21    7884   4595 6695
Complete

Read Sequence:sof-miR408b (21 nt)
Read Sequence:JN377537.1 Sugarcane bacilliform virus isolate BRU, complete
genome(7884 nt)
=-=-=-=-=-=-=-=-=-=-=-=-=-=-=-=-=-=-=-=-=-=-=-=-=-=-=-=-=-=-=-=-=
Performing Scan: sof-miR408b vs JN377537.1
=-=-=-=-=-=-=-=-=-=-=-=-=-=-=-=-=-=-=-=-=-=-=-=-=-=-=-=-=-=-=-=-=

   Forward: Score: 154.000000  Q:2 to 15  R:4595 to 4615 Align Len (13) (84.62%)
(84.62%)

   Query:    3' cggucccUUCUCCGUCACGUc 5'
                       |||  ||||||||
   Ref:      5' acaaaccAAGCAGCAGTGCAg 3'

   Energy:  -21.350000 kCal/Mol

Scores for this hit:
>sof-miR408b      JN377537.1  154.00      -21.35      2 15  4595 4615   13
      84.62%      84.62%
```

```
   Forward: Score: 135.000000  Q:2 to 16  R:6695 to 6715 Align Len (14) (71.43%)
(78.57%)

   Query:    3' cgguccCUUCUCCGUCACGUc 5'
                   |||  | :||||||
   Ref:      5' ctgtcaGAACTGATAGTGCAg 3'

   Energy:  -19.190001 kCal/Mol

Scores for this hit:
>sof-miR408b      JN377537.1  135.00      -19.19      2 16  6695 6715   14
      71.43%      78.57%

Score for this Scan:
Seq1,Seq2,Tot Score,Tot Energy,Max Score,Max Energy,Strand,Len1,Len2,Positions
>>sof-miR408b     JN377537.1  289.00      -40.54      154.00      -21.35      13
      21    7884   4595 6695
Complete

Read Sequence:sof-miR408c (21 nt)
Read Sequence:JN377537.1 Sugarcane bacilliform virus isolate BRU, complete
genome(7884 nt)
=-=-=-=-=-=-=-=-=-=-=-=-=-=-=-=-=-=-=-=-=-=-=-=-=-=-=-=-=-=-=-=-=
Performing Scan: sof-miR408c vs JN377537.1
=-=-=-=-=-=-=-=-=-=-=-=-=-=-=-=-=-=-=-=-=-=-=-=-=-=-=-=-=-=-=-=-=

   Forward: Score: 154.000000  Q:2 to 15  R:4595 to 4615 Align Len (13) (84.62%)
(84.62%)

   Query:    3' cgguccCUUCUCCGUCACGUc 5'
                    |||  ||||||||
   Ref:      5' acaaaccAAGCAGCAGTGCAg 3'

   Energy:  -21.350000 kCal/Mol

Scores for this hit:
>sof-miR408c      JN377537.1  154.00      -21.35      2 15  4595 4615   13
      84.62%      84.62%


   Forward: Score: 135.000000  Q:2 to 16  R:6695 to 6715 Align Len (14) (71.43%)
(78.57%)

   Query:    3' cgguccCUUCUCCGUCACGUc 5'
                   |||  | :||||||
   Ref:      5' ctgtcaGAACTGATAGTGCAg 3'

   Energy:  -19.190001 kCal/Mol

Scores for this hit:
>sof-miR408c      JN377537.1  135.00      -19.19      2 16  6695 6715   14
      71.43%      78.57%

Score for this Scan:
Seq1,Seq2,Tot Score,Tot Energy,Max Score,Max Energy,Strand,Len1,Len2,Positions
>>sof-miR408c     JN377537.1  289.00      -40.54      154.00      -21.35      14
      21    7884   4595 6695
Complete

Read Sequence:sof-miR408d (21 nt)
Read Sequence:JN377537.1 Sugarcane bacilliform virus isolate BRU, complete
genome(7884 nt)
=-=-=-=-=-=-=-=-=-=-=-=-=-=-=-=-=-=-=-=-=-=-=-=-=-=-=-=-=-=-=-=-=
```

```
Performing Scan: sof-miR408d vs JN377537.1
=-=-=-=-=-=-=-=-=-=-=-=-=-=-=-=-=-=-=-=-=-=-=-=-=-=-=-=-=-=-=-=

   Forward: Score: 154.000000  Q:2 to 15  R:4595 to 4615 Align Len (13) (84.62%)
(84.62%)

   Query:    3' cggucccUUCUCCGUCACGUc 5'
                       |||  ||||||||
   Ref:      5' acaaaccAAGCAGCAGTGCAg 3'

   Energy:  -21.350000 kCal/Mol

Scores for this hit:
>sof-miR408d      JN377537.1  154.00      -21.35      2 15  4595 4615   13
      84.62%      84.62%


   Forward: Score: 135.000000  Q:2 to 16  R:6695 to 6715 Align Len (14) (71.43%)
(78.57%)

   Query:    3' cgguccCUUCUCCGUCACGUc 5'
                      |||  | :||||||
   Ref:      5' ctgtcaGAACTGATAGTGCAg 3'

   Energy:  -19.190001 kCal/Mol

Scores for this hit:
>sof-miR408d      JN377537.1  135.00      -19.19      2 16  6695 6715   14
      71.43%      78.57%

Score for this Scan:
Seq1,Seq2,Tot Score,Tot Energy,Max Score,Max Energy,Strand,Len1,Len2,Positions
>>sof-miR408d     JN377537.1  289.00      -40.54      154.00      -21.35      15
      21    7884   4595 6695
Complete

Read Sequence:sof-miR408e (21 nt)
Read Sequence:JN377537.1 Sugarcane bacilliform virus isolate BRU, complete
genome(7884 nt)
=-=-=-=-=-=-=-=-=-=-=-=-=-=-=-=-=-=-=-=-=-=-=-=-=-=-=-=-=-=-=-=
Performing Scan: sof-miR408e vs JN377537.1
=-=-=-=-=-=-=-=-=-=-=-=-=-=-=-=-=-=-=-=-=-=-=-=-=-=-=-=-=-=-=-=

   Forward: Score: 147.000000  Q:2 to 15  R:4595 to 4615 Align Len (14) (85.71%)
(85.71%)

   Query:    3' cggucccUUC-UCAGUCACGUc 5'
                       ||| || |||||||
   Ref:      5' acaaaccAAGCAG-CAGTGCAg 3'

   Energy:  -19.530001 kCal/Mol

Scores for this hit:
>sof-miR408e      JN377537.1  147.00      -19.53      2 15  4595 4615   14
      85.71%      85.71%


   Forward: Score: 142.000000  Q:2 to 15  R:242 to 262 Align Len (13) (76.92%)
(92.31%)

   Query:    3' cggucccUUCUCAGUCACGUc 5'
                       :| |||:||||||
   Ref:      5' taacgttGATAGTTAGTGCAa 3'
```

```
   Energy:  -17.010000 kCal/Mol

Scores for this hit:
>sof-miR408e      JN377537.1  142.00      -17.01      2 15  242 262     13
      76.92%      92.31%


   Forward: Score: 135.000000  Q:2 to 16  R:6695 to 6715 Align Len (14) (71.43%)
(78.57%)

   Query:    3' cgguccCUUCUCAGUCACGUc 5'
                      |||  | :||||||
   Ref:      5' ctgtcaGAACTGATAGTGCAg 3'

   Energy:  -19.190001 kCal/Mol

Scores for this hit:
>sof-miR408e      JN377537.1  135.00      -19.19      2 16  6695 6715   14
      71.43%      78.57%

Score for this Scan:
Seq1,Seq2,Tot Score,Tot Energy,Max Score,Max Energy,Strand,Len1,Len2,Positions
>>sof-miR408e     JN377537.1  424.00      -55.73      147.00      -19.53      16
      21    7884   4595 242 6695
Complete

Read Sequence:ssp-miR166 (21 nt)
Read Sequence:JN377537.1 Sugarcane bacilliform virus isolate BRU, complete
genome(7884 nt)
=-=-=-=-=-=-=-=-=-=-=-=-=-=-=-=-=-=-=-=-=-=-=-=-=-=-=-=-=-=-=-=-=-=-=-=-=
Performing Scan: ssp-miR166 vs JN377537.1
=-=-=-=-=-=-=-=-=-=-=-=-=-=-=-=-=-=-=-=-=-=-=-=-=-=-=-=-=-=-=-=-=-=-=-=-=

   Forward: Score: 136.000000  Q:2 to 17  R:1986 to 2006 Align Len (15) (80.00%)
(86.67%)

   Query:    3' ccccuUACUUCGGACCAGGCu 5'
                     ||| |||:||||| |
   Ref:      5' actacATGCAGCTTGGTCGGg 3'

   Energy:  -20.959999 kCal/Mol

Scores for this hit:
>ssp-miR166 JN377537.1  136.00      -20.96      2 17  1986 2006   15    80.00%
      86.67%


   Forward: Score: 134.000000  Q:2 to 20  R:1449 to 1470 Align Len (19) (73.68%)
(78.95%)

   Query:    3' ccCCUUACUUC-GGACCAGGCu 5'
                  ||| |||:| |  ||||||
   Ref:      5' aaGGACTGAGGACAGGGTCCGa 3'

   Energy:  -27.850000 kCal/Mol

Scores for this hit:
>ssp-miR166 JN377537.1  134.00      -27.85      2 20  1449 1470   19    73.68%
      78.95%

Score for this Scan:
Seq1,Seq2,Tot Score,Tot Energy,Max Score,Max Energy,Strand,Len1,Len2,Positions
>>ssp-miR166      JN377537.1  270.00      -48.81      136.00      -27.85      17
      21    7884   1986 1449
```

```
Complete

Read Sequence:ssp-miR169 (21 nt)
Read Sequence:JN377537.1 Sugarcane bacilliform virus isolate BRU, complete
genome(7884 nt)
=-=-=-=-=-=-=-=-=-=-=-=-=-=-=-=-=-=-=-=-=-=-=-=-=-=-=-=-=-=-=-=-=
Performing Scan: ssp-miR169 vs JN377537.1
=-=-=-=-=-=-=-=-=-=-=-=-=-=-=-=-=-=-=-=-=-=-=-=-=-=-=-=-=-=-=-=-=

   Forward: Score: 145.000000  Q:2 to 18  R:7748 to 7768 Align Len (16) (68.75%)
(87.50%)

   Query:    3' ggccGUUCAGUAGGAACCGAu 5'
                    ::|||||  :||||||
   Ref:      5' tattTGAGTCAAGTTTGGCTt 3'

   Energy:  -17.389999 kCal/Mol

Scores for this hit:
>ssp-miR169 JN377537.1  145.00      -17.39      2 18  7748 7768   16    68.75%
      87.50%

Score for this Scan:
Seq1,Seq2,Tot Score,Tot Energy,Max Score,Max Energy,Strand,Len1,Len2,Positions
>>ssp-miR169      JN377537.1  145.00      -17.39      145.00      -17.39      18
      21    7884   7748
Complete

Read Sequence:ssp-miR437a (21 nt)
Read Sequence:JN377537.1 Sugarcane bacilliform virus isolate BRU, complete
genome(7884 nt)
=-=-=-=-=-=-=-=-=-=-=-=-=-=-=-=-=-=-=-=-=-=-=-=-=-=-=-=-=-=-=-=-=
Performing Scan: ssp-miR437a vs JN377537.1
=-=-=-=-=-=-=-=-=-=-=-=-=-=-=-=-=-=-=-=-=-=-=-=-=-=-=-=-=-=-=-=-=
Score for this Scan:
No Hits Found above Threshold
Complete

Read Sequence:ssp-miR437b (21 nt)
Read Sequence:JN377537.1 Sugarcane bacilliform virus isolate BRU, complete
genome(7884 nt)
=-=-=-=-=-=-=-=-=-=-=-=-=-=-=-=-=-=-=-=-=-=-=-=-=-=-=-=-=-=-=-=-=
Performing Scan: ssp-miR437b vs JN377537.1
=-=-=-=-=-=-=-=-=-=-=-=-=-=-=-=-=-=-=-=-=-=-=-=-=-=-=-=-=-=-=-=-=
Score for this Scan:
No Hits Found above Threshold
Complete

Read Sequence:ssp-miR437c (21 nt)
Read Sequence:JN377537.1 Sugarcane bacilliform virus isolate BRU, complete
genome(7884 nt)
=-=-=-=-=-=-=-=-=-=-=-=-=-=-=-=-=-=-=-=-=-=-=-=-=-=-=-=-=-=-=-=-=
Performing Scan: ssp-miR437c vs JN377537.1
=-=-=-=-=-=-=-=-=-=-=-=-=-=-=-=-=-=-=-=-=-=-=-=-=-=-=-=-=-=-=-=-=
Score for this Scan:
No Hits Found above Threshold
Complete

Read Sequence:ssp-miR528 (21 nt)
Read Sequence:JN377537.1 Sugarcane bacilliform virus isolate BRU, complete
genome(7884 nt)
=-=-=-=-=-=-=-=-=-=-=-=-=-=-=-=-=-=-=-=-=-=-=-=-=-=-=-=-=-=-=-=-=
Performing Scan: ssp-miR528 vs JN377537.1
=-=-=-=-=-=-=-=-=-=-=-=-=-=-=-=-=-=-=-=-=-=-=-=-=-=-=-=-=-=-=-=-=
```

```
Score for this Scan:
No Hits Found above Threshold
Complete

Read Sequence:ssp-miR827 (21 nt)
Read Sequence:JN377537.1 Sugarcane bacilliform virus isolate BRU, complete
genome(7884 nt)
=-=-=-=-=-=-=-=-=-=-=-=-=-=-=-=-=-=-=-=-=-=-=-=-=-=-=-=-=-=-=-=-=
Performing Scan: ssp-miR827 vs JN377537.1
=-=-=-=-=-=-=-=-=-=-=-=-=-=-=-=-=-=-=-=-=-=-=-=-=-=-=-=-=-=-=-=-=

   Forward: Score: 142.000000  Q:2 to 16  R:2816 to 2837 Align Len (15) (80.00%)
(86.67%)

   Query:    3' acaaacGAC-UACCAGUAGAUu 5'
                      ||| | ||||||||:
   Ref:      5' ctacatCTGAAGGGTCATCTGc 3'

   Energy:  -19.730000 kCal/Mol

Scores for this hit:
>ssp-miR827 JN377537.1  142.00      -19.73      2 16  2816 2837   15    80.00%
      86.67%

Score for this Scan:
Seq1,Seq2,Tot Score,Tot Energy,Max Score,Max Energy,Strand,Len1,Len2,Positions
>>ssp-miR827      JN377537.1  142.00      -19.73      142.00      -19.73      23
      21    7884   2816
Complete

Read Sequence:ssp-miR444a (21 nt)
Read Sequence:JN377537.1 Sugarcane bacilliform virus isolate BRU, complete
genome(7884 nt)
=-=-=-=-=-=-=-=-=-=-=-=-=-=-=-=-=-=-=-=-=-=-=-=-=-=-=-=-=-=-=-=-=
Performing Scan: ssp-miR444a vs JN377537.1
=-=-=-=-=-=-=-=-=-=-=-=-=-=-=-=-=-=-=-=-=-=-=-=-=-=-=-=-=-=-=-=-=

   Forward: Score: 136.000000  Q:3 to 15  R:3293 to 3312 Align Len (12) (91.67%)
(91.67%)

   Query:    3' uucgaacUCCGUUGUUGACgu 5'
                       ||| ||||||||
   Ref:      5' ctgaacaAGG-AACAACTGga 3'

   Energy:  -15.230000 kCal/Mol

Scores for this hit:
>ssp-miR444a      JN377537.1  136.00      -15.23      3 15  3293 3312   12
      91.67%      91.67%


   Forward: Score: 135.000000  Q:3 to 18  R:1679 to 1701 Align Len (17) (76.47%)
(82.35%)

   Query:    3' uucgAACUC-CGU-UGUUGACgu 5'
                    |||:| | | |||||||
   Ref:      5' ttagTTGGGTGAAGACAACTGat 3'

   Energy:  -16.250000 kCal/Mol

Scores for this hit:
>ssp-miR444a      JN377537.1  135.00      -16.25      3 18  1679 1701   17
      76.47%      82.35%
```

```
   Forward: Score: 134.000000  Q:3 to 19  R:6184 to 6204 Align Len (16) (75.00%)
(87.50%)

   Query:    3' uucGAACUCCGUUGUUGACgu 5'
                   ||||:  ||||||:||
   Ref:      5' tttCTTGGTTCAACAATTGgt 3'

   Energy:  -18.040001 kCal/Mol

Scores for this hit:
>ssp-miR444a      JN377537.1  134.00      -18.04      3 19  6184 6204   16
      75.00%      87.50%


   Forward: Score: 132.000000  Q:2 to 18  R:1301 to 1326 Align Len (21) (61.90%)
(71.43%)

   Query:    3' uucgAACUCCGU-----UGUUGACGu 5'
                    ||||| :|     |||:||||
   Ref:      5' gttaTTGAGTTACTTCTACAGCTGCa 3'

   Energy:  -17.440001 kCal/Mol

Scores for this hit:
>ssp-miR444a      JN377537.1  132.00      -17.44      2 18  1301 1326   21
      61.90%      71.43%

Score for this Scan:
Seq1,Seq2,Tot Score,Tot Energy,Max Score,Max Energy,Strand,Len1,Len2,Positions
>>ssp-miR444a     JN377537.1  537.00      -66.96      136.00      -18.04      24
      21    7884   3293 1679 6184 1301
Complete

Read Sequence:ssp-miR444b (21 nt)
Read Sequence:JN377537.1 Sugarcane bacilliform virus isolate BRU, complete
genome(7884 nt)
=-=-=-=-=-=-=-=-=-=-=-=-=-=-=-=-=-=-=-=-=-=-=-=-=-=-=-=-=-=-=-=
Performing Scan: ssp-miR444b vs JN377537.1
=-=-=-=-=-=-=-=-=-=-=-=-=-=-=-=-=-=-=-=-=-=-=-=-=-=-=-=-=-=-=-=

   Forward: Score: 136.000000  Q:3 to 15  R:3293 to 3312 Align Len (12) (91.67%)
(91.67%)

   Query:    3' uucgaacUCCGUUGUUGACgu 5'
                       ||| ||||||||
   Ref:      5' ctgaacaAGG-AACAACTGga 3'

   Energy:  -15.230000 kCal/Mol

Scores for this hit:
>ssp-miR444b      JN377537.1  136.00      -15.23      3 15  3293 3312   12
      91.67%      91.67%


   Forward: Score: 135.000000  Q:3 to 18  R:1679 to 1701 Align Len (17) (76.47%)
(82.35%)

   Query:    3' uucgAACUC-CGU-UGUUGACgu 5'
                    |||:| | | |||||||
   Ref:      5' ttagTTGGGTGAAGACAACTGat 3'

   Energy:  -16.250000 kCal/Mol
```

```
Scores for this hit:
>ssp-miR444b      JN377537.1  135.00      -16.25      3 18  1679 1701   17
      76.47%      82.35%


   Forward: Score: 134.000000  Q:3 to 19  R:6184 to 6204 Align Len (16) (75.00%)
(87.50%)

   Query:    3' uucGAACUCCGUUGUUGACgu 5'
                   ||||:  ||||||:||
   Ref:      5' tttCTTGGTTCAACAATTGgt 3'

   Energy:  -18.040001 kCal/Mol

Scores for this hit:
>ssp-miR444b      JN377537.1  134.00      -18.04      3 19  6184 6204   16
      75.00%      87.50%


   Forward: Score: 132.000000  Q:2 to 18  R:1301 to 1326 Align Len (21) (61.90%)
(71.43%)

   Query:    3' uucgAACUCCGU-----UGUUGACGu 5'
                    ||||| :|     |||:||||
   Ref:      5' gttaTTGAGTTACTTCTACAGCTGCa 3'

   Energy:  -17.440001 kCal/Mol

Scores for this hit:
>ssp-miR444b      JN377537.1  132.00      -17.44      2 18  1301 1326   21
      61.90%      71.43%

Score for this Scan:
Seq1,Seq2,Tot Score,Tot Energy,Max Score,Max Energy,Strand,Len1,Len2,Positions
>>ssp-miR444b     JN377537.1  537.00      -66.96      136.00      -18.04      25
      21    7884   3293 1679 6184 1301
Complete

Read Sequence:ssp-miR444c-3p (21 nt)
Read Sequence:JN377537.1 Sugarcane bacilliform virus isolate BRU, complete
genome(7884 nt)
=-=-=-=-=-=-=-=-=-=-=-=-=-=-=-=-=-=-=-=-=-=-=-=-=-=-=-=-=-=-=-=-=
Performing Scan: ssp-miR444c-3p vs JN377537.1
=-=-=-=-=-=-=-=-=-=-=-=-=-=-=-=-=-=-=-=-=-=-=-=-=-=-=-=-=-=-=-=-=

   Forward: Score: 138.000000  Q:3 to 20  R:1680 to 1701 Align Len (18) (66.67%)
(83.33%)

   Query:    3' uuCGAACU-CUGUUGUUGACgu 5'
                  |:| |: || :|||||||
   Ref:      5' taGTTGGGTGAAGACAACTGat 3'

   Energy:  -16.969999 kCal/Mol

Scores for this hit:
>ssp-miR444c-3p   JN377537.1  138.00      -16.97      3 20  1680 1701   18
      66.67%      83.33%


   Forward: Score: 134.000000  Q:3 to 19  R:6184 to 6204 Align Len (16) (75.00%)
(87.50%)

   Query:    3' uucGAACUCUGUUGUUGACgu 5'
                   ||||:  ||||||:||
```

```
   Ref:      5' tttCTTGGTTCAACAATTGgt 3'

   Energy:  -18.950001 kCal/Mol

Scores for this hit:
>ssp-miR444c-3p   JN377537.1  134.00      -18.95      3 19  6184 6204   16
      75.00%      87.50%


   Forward: Score: 132.000000  Q:2 to 19  R:328 to 354 Align Len (23) (65.22%)
(69.57%)

   Query:    3' uucGAACUCU-GUU-----GUUGACGu 5'
                   || |||| |||     |||:|||
   Ref:      5' aacCTAGAGATCAAAAGTTCAATTGCa 3'

   Energy:  -16.080000 kCal/Mol

Scores for this hit:
>ssp-miR444c-3p   JN377537.1  132.00      -16.08      2 19  328 354     23
      65.22%      69.57%


   Forward: Score: 132.000000  Q:2 to 18  R:1301 to 1326 Align Len (21) (61.90%)
(71.43%)

   Query:    3' uucgAACUCUGU-----UGUUGACGu 5'
                    ||||| :|     |||:||||
   Ref:      5' gttaTTGAGTTACTTCTACAGCTGCa 3'

   Energy:  -18.590000 kCal/Mol

Scores for this hit:
>ssp-miR444c-3p   JN377537.1  132.00      -18.59      2 18  1301 1326   21
      61.90%      71.43%

Score for this Scan:
Seq1,Seq2,Tot Score,Tot Energy,Max Score,Max Energy,Strand,Len1,Len2,Positions
>>ssp-miR444c-3p  JN377537.1  536.00      -70.59      138.00      -18.95      26
      21    7884   1680 6184 328 1301
Complete

Read Sequence:ssp-miR1128 (21 nt)
Read Sequence:JN377537.1 Sugarcane bacilliform virus isolate BRU, complete
genome(7884 nt)
=-=-=-=-=-=-=-=-=-=-=-=-=-=-=-=-=-=-=-=-=-=-=-=-=-=-=-=-=-=-=-=-=
Performing Scan: ssp-miR1128 vs JN377537.1
=-=-=-=-=-=-=-=-=-=-=-=-=-=-=-=-=-=-=-=-=-=-=-=-=-=-=-=-=-=-=-=-=
Score for this Scan:
No Hits Found above Threshold
Complete

Read Sequence:ssp-miR1432 (21 nt)
Read Sequence:JN377537.1 Sugarcane bacilliform virus isolate BRU, complete
genome(7884 nt)
=-=-=-=-=-=-=-=-=-=-=-=-=-=-=-=-=-=-=-=-=-=-=-=-=-=-=-=-=-=-=-=-=
Performing Scan: ssp-miR1432 vs JN377537.1
=-=-=-=-=-=-=-=-=-=-=-=-=-=-=-=-=-=-=-=-=-=-=-=-=-=-=-=-=-=-=-=-=
Score for this Scan:
No Hits Found above Threshold
Complete

Scan Complete
```

Results from psRNATarget

miRNA_Acc. Target_Acc. Expectation UPE miRNA_start miRNA_end Target_start
Target_end miRNA_aligned_fragment alignment Target_aligned_fragment
Inhibition Target_Desc. Multiplicity

sof-miR159a Your_Sequence 7.0 -1.0 1 21 5108 5127
UUUGGAUUGAAGGGAGCUCUG ::: :.:.: ::: :..:: AUGAG-UUCUUGCAAGCUGAA
Translation 4

sof-miR159b Your_Sequence 7.0 -1.0 1 21 5108 5127
UUUGGAUUGAAGGGAGCUCUG ::: :.:.: ::: :..:: AUGAG-UUCUUGCAAGCUGAA
Translation 4

sof-miR159d Your_Sequence 7.0 -1.0 1 21 5108 5127
UUUGGAUUGAAGGGAGCUCUG ::: :.:.: ::: :..:: AUGAG-UUCUUGCAAGCUGAA
Translation 4

sof-miR159e Your_Sequence 7.0 -1.0 1 21 5534 5552
UUUGGAUUGAAAGGAGCUCUU :::::  : ::::::::.: UUGAGCU--UAUCAAUCCAGA
Translation 7

sof-miR159e Your_Sequence 7.0 -1.0 1 21 5576 5596
UUUGGAUUGAAAGGAGCUCUU : ::: :::::::.: UUACAAUGCUUACAAUCCAGA
Translation 7

sof-miR408e Your_Sequence 7.25 -1.0 1 21 174 196 CUGCACUGAC--
UCUUCCCUGGC :.:::.:::  ::.: :::.: UCUAGGGGAGAACGUUAAUGCGG Translation
5

sof-miR159a Your_Sequence 7.5 -1.0 1 21 5576 5596
UUUGGAUUGAAGGGAGCUCUG : :.: :::::::.: UUACAAUGCUUACAAUCCAGA
Translation 4

sof-miR159a Your_Sequence 7.5 -1.0 1 21 2825 2845
UUUGGAUUGAAGGGAGCUCUG ::.:.  .:: :::::... AAGGGUCAUCUGCAAUCUGGC
Translation 4

sof-miR159b Your_Sequence 7.5 -1.0 1 21 5576 5596
UUUGGAUUGAAGGGAGCUCUG : :.: :::::::.: UUACAAUGCUUACAAUCCAGA
Translation 4

sof-miR159b Your_Sequence 7.5 -1.0 1 21 2825 2845
UUUGGAUUGAAGGGAGCUCUG ::.:.  .:: :::::... AAGGGUCAUCUGCAAUCUGGC
Translation 4

sof-miR159c Your_Sequence 7.5 -1.0 1 21 2825 2845
CUUGGAUUGAAGGGAGCUCCU : :.:.  .:: :::::... AAGGGUCAUCUGCAAUCUGGC
Translation 4

sof-miR159d Your_Sequence 7.5 -1.0 1 21 5576 5596
UUUGGAUUGAAGGGAGCUCUG : :.: :::::::.: UUACAAUGCUUACAAUCCAGA
Translation 4

sof-miR159d Your_Sequence 7.5 -1.0 1 21 2825 2845
UUUGGAUUGAAGGGAGCUCUG ::.:. .:: :::::... AAGGGUCAUCUGCAAUCUGGC
Translation 4

sof-miR168a Your_Sequence 7.5 -1.0 1 21 6792 6811
UCGCUUGGUGCAGAUCGGGAC :. :: ::: .:::: ::: CACUGGAGCUGGGCCAA-CGA
Translation 3

sof-miR408e Your_Sequence 7.5 -1.0 1 21 5293 5313
CUGCACUGACUCUUCCCUGGC :::.::::::: ..: UCGCUGGAGGAGUCAGAUUGG
Cleavage 5

sof-miR156 Your_Sequence 8.0 -1.0 1 20 818 837
UGACAGAAGAGAGUGAGCAC ..: :.::::::: AGCAGAGUUGUUUCUGUCA
Translation 3

sof-miR156 Your_Sequence 8.0 -1.0 1 20 7609 7628 UGACAGAAG-AGAGUGAGCAC :::: ::::.: : :.::::. GUGC-CACUUUACCUUUGUCG Cleavage
3

sof-miR408a Your_Sequence 8.0 -1.0 1 21 4595 4615
CUGCACUGCCUCUUCCCUGGC ::: ::::::::: ACAAACCAAGCAGCAGUGCAG
Translation 3

sof-miR408a Your_Sequence 8.0 -1.0 1 21 3669 3689
CUGCACUGCCUCUUCCCUGGC :: ::::::.: :.: AUAUGGAAAGAGGUAUGGUAC
Cleavage 3

sof-miR408a Your_Sequence 8.0 -1.0 1 21 1766 1786
CUGCACUGCCUCUUCCCUGGC :: .:::: .:: ::: UGAAGAAGAGAGCUAGGGCAU
Cleavage 3

sof-miR408b Your_Sequence 8.0 -1.0 1 21 4595 4615
CUGCACUGCCUCUUCCCUGGC ::: ::::::::: ACAAACCAAGCAGCAGUGCAG
Translation 3

sof-miR408b Your_Sequence 8.0 -1.0 1 21 3669 3689
CUGCACUGCCUCUUCCCUGGC :: ::::::.: :.: AUAUGGAAAGAGGUAUGGUAC
Cleavage 3

sof-miR408b Your_Sequence 8.0 -1.0 1 21 1766 1786
CUGCACUGCCUCUUCCCUGGC :: .:::: .:: ::: UGAAGAAGAGAGCUAGGGCAU
Cleavage 3

sof-miR408c Your_Sequence 8.0 -1.0 1 21 4595 4615
CUGCACUGCCUCUUCCCUGGC ::: ::::::::: ACAAACCAAGCAGCAGUGCAG
Translation 3

sof-miR408c Your_Sequence 8.0 -1.0 1 21 3669 3689
CUGCACUGCCUCUUCCCUGGC :: ::::::.: :.: AUAUGGAAAGAGGUAUGGUAC
Cleavage 3

sof-miR408c Your_Sequence 8.0 -1.0 1 21 1766 1786
CUGCACUGCCUCUUCCCUGGC :: .:::: .:: ::: UGAAGAAGAGAGCUAGGGCAU
Cleavage 3

sof-miR408d Your_Sequence 8.0 -1.0 1 21 4595 4615
CUGCACUGCCUCUUCCCUGGC ::: :::::::: ACAAACCAAGCAGCAGUGCAG
Translation 3

sof-miR408d Your_Sequence 8.0 -1.0 1 21 3669 3689
CUGCACUGCCUCUUCCCUGGC :: ::::::.: :.: AUAUGGAAAGAGGUAUGGUAC
Cleavage 3

sof-miR408d Your_Sequence 8.0 -1.0 1 21 1766 1786
CUGCACUGCCUCUUCCCUGGC :: .:::: .:: ::: UGAAGAAGAGAGCUAGGGCAU
Cleavage 3

sof-miR408e Your_Sequence 8.0 -1.0 1 21 1766 1786
CUGCACUGACUCUUCCCUGGC :: .:::: .:: ::: UGAAGAAGAGAGCUAGGGCAU
Cleavage 5

sof-miR408e Your_Sequence 8.0 -1.0 1 21 5683 5702
CUGCACUGACUCUUCCCUGGC ..: : .:::::: ::: UUUAUUGUGGAGUCAG-GCAC
Cleavage 5

sof-miR159e Your_Sequence 8.25 -1.0 1 21 820 842 UUUG--
GAUUGAAAGGAGCUCUU ::::.: .:::: .:: ::.: CAGAGUUGUUUUCUGUCACCAGACleavage
7

sof-miR396 Your_Sequence 8.25 -1.0 1 21 5368 5390 UUCCACAGCUUU--
CUUGAACUG : :::: .:::.:: :::: UCUUGCAAGCAGAAGUUGAGGAA Cleavage
2

sof-miR156 Your_Sequence 8.5 -1.0 1 20 567 586
UGACAGAAGAGAGUGAGCAC : :: ::: ::::.: GUAAACCCUAUCUCCUGUUA
Translation 3

sof-miR159a Your_Sequence 8.5 -1.0 1 21 6609 6629
UUUGGAUUGAAGGGAGCUCUG :: :: ::: ..:: ::.: CAAAGGAACCUGUGAUGCAGA
Translation 4

sof-miR159b Your_Sequence 8.5 -1.0 1 21 6609 6629
UUUGGAUUGAAGGGAGCUCUG :: :: ::: ..:: ::.: CAAAGGAACCUGUGAUGCAGA
Translation 4

sof-miR159c Your_Sequence 8.5 -1.0 1 21 6177 6197
CUUGGAUUGAAGGGAGCUCCU :: .: :... :...:.::: AGUGGAUUUUCUUGGUUCAAC
Translation 4

sof-miR159c Your_Sequence 8.5 -1.0 1 21 5576 5596
CUUGGAUUGAAGGGAGCUCCU : :.: :::::::. UUACAAUGCUUACAAUCCAGA
Translation 4

sof-miR159c	Your_Sequence 8.5	-1.0	1	21	1003	1022
CUUGGAUUGAAGGGAGCUCCU	:: .:  ::::.: ::.:::	AGCGGAGGCCUUUA-UCUAAG
Cleavage	4

sof-miR159d	Your_Sequence 8.5	-1.0	1	21	6609	6629
UUUGGAUUGAAGGGAGCUCUG	:: ::  ::: ..:: ::.:	CAAAGGAACCUGUGAUGCAGA
Translation	4

sof-miR159e	Your_Sequence 8.5	-1.0	1	21	2647	2667
UUUGGAUUGAAAGGAGCUCUU	:.:::: :::...:	AUUUCAACUUUUCUAUCUGGA
Cleavage	7

sof-miR159e	Your_Sequence 8.5	-1.0	1	21	713	733
UUUGGAUUGAAAGGAGCUCUU	::  ::: .::..:::	AAUUUCUAAUUUAGAUUUAAA
Cleavage	7

sof-miR159e	Your_Sequence 8.5	-1.0	1	21	4309	4329
UUUGGAUUGAAAGGAGCUCUU	: : .: :::. .::::	CUAUGGUGUUAUCAGAUCAAA
Translation	7

sof-miR159e	Your_Sequence 8.5	-1.0	1	21	6661	6681
UUUGGAUUGAAAGGAGCUCUU	: :::  .::::.:  ...:	AUGAGACUCUUUUAUCUUGAU
Cleavage	7

sof-miR168a	Your_Sequence 8.5	-1.0	1	21	3451	3471
UCGCUUGGUGCAGAUCGGGAC	::: :::::::. :	AAUGAAUUCUUCACCAAGUUA
Translation	3

sof-miR168a	Your_Sequence 8.5	-1.0	1	21	4046	4066
UCGCUUGGUGCAGAUCGGGAC	:: .::.:: ::: :.	GGGAAGAAUUGUACAAAGAGG
Cleavage	3

sof-miR396	Your_Sequence 8.5	-1.0	1	21	79	104	UUCCACAGCU-----
UUCUUGAACUG	::::.:::.:	::::::: ::	GAGUUUAAGGACAACUAGCUGUGCAA
Translation	2

sof-miR408e	Your_Sequence 8.5	-1.0	1	21	242	262
CUGCACUGACUCUUCCCUGGC	.: :::.::::::	UAACGUUGAUAGUUAGUGCAA
Cleavage	5

#Please import the downloaded file into Microsoft Excel or other speadsheet software

miRNA_Acc.	Target_Acc.	Expectation	UPE$	miRNA_start	miRNA_end	Target_start
Target_end	miRNA_aligned_fragment	alignment	Target_aligned_fragment
Inhibition	Target_Desc.	Multiplicity

ssp-miR166	Your_Sequence 6.0	-1.0	1	21	7750	7770
UCGGACCAGGCUUCAUUCCCC	::.: :::..::: ..::	UUUGAGUCAAGUUUGGCUUGA
Cleavage	5

ssp-miR437c	Your_Sequence 6.0	-1.0	1	21	2974	2994
AAAGUUAGAGAAGUCUGACUU	.: ::::: ::::.:::. :	GAAUCAGAGUUCUUUAAUCUA
Cleavage	5

ssp-miR444a Your_Sequence 6.0 -1.0 1 21 6797 6816
UGCAGUUGUUGCCUCAAGCUU .:::: :.: ::::.:.::.. GAGCU-GGGCCAACGAUUGUG
Cleavage 2

ssp-miR444b.1 Your_Sequence 6.0 -1.0 1 21 7079 7099
UUGUGGCUUUCUUGCAAGUUG ::::.:::: :..::: GUGGAUGCAGGAAACCUGCAA
Cleavage 16

ssp-miR444b.2 Your_Sequence 6.0 -1.0 1 21 6797 6816
UGCAGUUGUUGCCUCAAGCUU .:::: :.: ::::.:.::.. GAGCU-GGGCCAACGAUUGUG
Cleavage 2

ssp-miR444c-3p Your_Sequence 6.0 -1.0 1 21 6797 6816
UGCAGUUGUUGUCUCAAGCUU .:::: :.: ::::.:.::.. GAGCU-GGGCCAACGAUUGUG
Cleavage 8

ssp-miR437a Your_Sequence 6.5 -1.0 1 21 2974 2994
AAAGUUAGAGAAGUUUGACUU .: ::::.: ::::.:::. : GAAUCAGAGUUCUUUAAUCUA
Cleavage 5

ssp-miR444a Your_Sequence 6.5 -1.0 1 21 6865 6885
UGCAGUUGUUGCCUCAAGCUU :.:::  :::::: : .::: AGGCUCAAGGCAAAACUUGCU
Cleavage 2

ssp-miR444b.1 Your_Sequence 6.5 -1.0 1 21 6822 6842
UUGUGGCUUUCUUGCAAGUUG :: :  :::.::  ::::: GCACAUCAAAGGAAAACACAA
Cleavage 16

ssp-miR444b.2 Your_Sequence 6.5 -1.0 1 21 6865 6885
UGCAGUUGUUGCCUCAAGCUU :.:::  :::::: : .::: AGGCUCAAGGCAAAACUUGCU
Cleavage 2

ssp-miR159a Your_Sequence 7.0 -1.0 1 21 5108 5127
UUUGGAUUGAAGGGAGCUCUG ::: :.:.: ::: :..::: AUGAG-UUCUUGCAAGCUGAA
Translation 4

ssp-miR166 Your_Sequence 7.0 -1.0 1 21 1986 2006
UCGGACCAGGCUUCAUUCCCC ::: :::.::::: :. ACUACAUGCAGCUUGGUCGGG
Cleavage 5

ssp-miR166 Your_Sequence 7.0 -1.0 1 21 3863 3882
UCGGACCAGGCUUCAUUCCCC ::.:::: .:::: .::: ACAGAGUGAA-UCUGGCUCGA
Translation 5

ssp-miR437a Your_Sequence 7.0 -1.0 1 21 2646 2666
AAAGUUAGAGAAGUUUGACUU : :. :::::.:::: :: UAUUUCAACUUUUCUAUCUGG
Cleavage 5

ssp-miR437a Your_Sequence 7.0 -1.0 1 21 705 725
AAAGUUAGAGAAGUUUGACUU ::::  :.:::::.:: GGAACAAAAAUUUCUAAUUUA
Cleavage 5

ssp-miR437c Your_Sequence 7.0 -1.0 1 21 6659 6679
AAAGUUAGAGAAGUCUGACUU :: : :::: .:.:: ::: AAAUGAGACUCUUUUAUCUUG
Translation 5

ssp-miR444b.1 Your_Sequence 7.0 -1.0 1 21 7008 7027
UUGUGGCUUUCUUGCAAGUUG ::: :: ::::::: .:.. CAAGUU-CAAGAAAGAUAUGU
Cleavage 16

ssp-miR444b.1 Your_Sequence 7.0 -1.0 1 21 2951 2971
UUGUGGCUUUCUUGCAAGUUG :: :: ::.:::. .:: .: GAAAUUCCAGGAAGUUCAAGA
Cleavage 16

ssp-miR444b.1 Your_Sequence 7.0 -1.0 1 21 7139 7159
UUGUGGCUUUCUUGCAAGUUG : .:: :::: . ::::..: GAUUUUACAAGUGCGCCAUGA
Translation 16

ssp-miR444b.1 Your_Sequence 7.0 -1.0 1 21 7386 7406
UUGUGGCUUUCUUGCAAGUUG :.. : ::::: ::::.: GAGUCUAGAAGAACGCCAUAC
Cleavage 16

ssp-miR444c-3p Your_Sequence 7.0 -1.0 1 21 5248 5269
UGCAGUUGUUGUCUCAA-GCUU :::: :::.:: :: ::.::: AAGCAUUGGGAAAAGAAUUGCU
Translation 8

ssp-miR444c-3p Your_Sequence 7.0 -1.0 1 21 6865 6885
UGCAGUUGUUGUCUCAAGCUU :.::: ::.::: : .::: AGGCUCAAGGCAAAACUUGCU
Cleavage 8

ssp-miR159a Your_Sequence 7.5 -1.0 1 21 5576 5596
UUUGGAUUGAAGGGAGCUCUG : :.: :::::::.: UUACAAUGCUUACAAUCCAGA
Translation 4

ssp-miR159a Your_Sequence 7.5 -1.0 1 21 2825 2845
UUUGGAUUGAAGGGAGCUCUG ::.:. .:: :::::... AAGGGUCAUCUGCAAUCUGGC
Translation 4

ssp-miR166 Your_Sequence 7.5 -1.0 1 21 2060 2083
UCGGACCAGGCUUCAU---UCCCC :::: :::.:: .:: ::.::
UGGGAGCUAUGGAGAUUGAUCUGA Translation 5

ssp-miR168a Your_Sequence 7.5 -1.0 1 21 6792 6811
UCGCUUGGUGCAGAUCGGGAC :. :: ::: .:::: ::: CACUGGAGCUGGGCCAA-CGA
Translation 3

ssp-miR169 Your_Sequence 7.5 -1.0 1 21 7748 7768
UAGCCAAGGAUGACUUGCCGG ..::::: .:::::: UAUUUGAGUCAAGUUUGGCUU
Translation 3

ssp-miR437a Your_Sequence 7.5 -1.0 1 21 6659 6679
AAAGUUAGAGAAGUUUGACUU :: : :.::: .:.:: ::: AAAUGAGACUCUUUUAUCUUG
Translation 5

ssp-miR437c	Your_Sequence	7.5	-1.0	1	21	363	383
AAAGUUAGAGAAGUCUGACUU	:::: .:: :::.. ::	CUAUCAGUUUUAUCUGGAUUG
Translation	5

ssp-miR437c	Your_Sequence	7.5	-1.0	1	21	2647	2666
AAAGUUAGAGAAGUCUGACUU	::: ::::.:::: ::	AUUUCA-ACUUUUCUAUCUGG
Cleavage	5

ssp-miR444b.1	Your_Sequence	7.5	-1.0	1	21	1089	1110	UUGUGGCUUUCUU-
GCAAGUUG	: :::: . .::::::.:: .:	CUACUUCUUGAGAAAGUCAAGA	Cleavage	16

ssp-miR444b.1	Your_Sequence	7.5	-1.0	1	21	4466	4486
UUGUGGCUUUCUUGCAAGUUG	..:: :  ::.::::.. .:	AGGCUAGACAGGAAGCUGAGA
Cleavage	16

ssp-miR827	Your_Sequence	7.5	-1.0	1	21	7337	7357
UUAGAUGACCAUCAGCAAACA	: : :::: ::::.: :	GGCCUACAGAUGAUCAUUUCA
Cleavage	3

ssp-miR1128	Your_Sequence	8.0	-1.0	1	21	1553	1573
UACUACUCCCUCCGUCCCAAA	:: . :::::: :.::	AACAGGUUCGAGGGACUGGUU
Cleavage	1

ssp-miR156	Your_Sequence	8.0	-1.0	1	21	7608	7628	UGACAGAAG-
AGAGUGAGCACA	::::: ::::.: : :.::::.	UGUGC-CACUUUACCUUUGUCG	Cleavage
3

ssp-miR156	Your_Sequence	8.0	-1.0	1	21	817	837
UGACAGAAGAGAGUGAGCACA	..: :.:::::::	GAGCAGAGUUGUUUUCUGUCA
Translation	3

ssp-miR169	Your_Sequence	8.0	-1.0	1	21	2091	2112	UAGC-
CAAGGAUGACUUGCCGG	::: :::::. :: ::::	AGCACAACUCAUCUAUGUGCUA	Cleavage
3

ssp-miR169	Your_Sequence	8.0	-1.0	1	21	7626	7647
UAGCCAAGGAUGAC-UUGCCGG	.:::: : ::.. :.::: .::	UCGGCCACGUUGCCUUUGCUUA
Translation	3

ssp-miR408a	Your_Sequence	8.0	-1.0	1	21	4595	4615
CUGCACUGCCUCUUCCCUGGC	:::  ::::::::	ACAAACCAAGCAGCAGUGCAG
Translation	3

ssp-miR408a	Your_Sequence	8.0	-1.0	1	21	3669	3689
CUGCACUGCCUCUUCCCUGGC	:: ::::::.:  :.:	AUAUGGAAAGAGGUAUGGUAC
Cleavage	3

ssp-miR408a	Your_Sequence	8.0	-1.0	1	21	1766	1786
CUGCACUGCCUCUUCCCUGGC	::  .:::: .:: :::	UGAAGAAGAGAGCUAGGGCAU
Cleavage	3

ssp-miR408d Your_Sequence 8.0 -1.0 1 21 4595 4615
CUGCACUGCCUCUUCCCUGGC ::: :::::::: ACAAACCAAGCAGCAGUGCAG
Translation 3

ssp-miR408d Your_Sequence 8.0 -1.0 1 21 3669 3689
CUGCACUGCCUCUUCCCUGGC :: ::::::.: :.: AUAUGGAAAGAGGUAUGGUAC
Cleavage 3

ssp-miR408d Your_Sequence 8.0 -1.0 1 21 1766 1786
CUGCACUGCCUCUUCCCUGGC :: .:::: .:: ::: UGAAGAAGAGAGCUAGGGCAU
Cleavage 3

ssp-miR437a Your_Sequence 8.0 -1.0 1 21 363 383
AAAGUUAGAGAAGUUUGACUU :::. .:: :::.. :: CUAUCAGUUUUAUCUGGAUUG
Translation 5

ssp-miR437b Your_Sequence 8.0 -1.0 1 21 363 383
AAAGUUAGACAAGUUUGACAU : :::. .:: :::.. :: CUAUCAGUUUUAUCUGGAUUG
Translation 2

ssp-miR437c Your_Sequence 8.0 -1.0 1 21 705 725
AAAGUUAGAGAAGUCUGACUU :: : :.:::::.:: GGAACAAAAAUUUCUAAUUUA
Cleavage 5

ssp-miR444b.1 Your_Sequence 8.0 -1.0 1 21 3113 3133
UUGUGGCUUUCUUGCAAGUUG : .:: ..: ::. ..:::: CUGCUGAUGACAAGAUUACAA
Translation 16

ssp-miR444c-3p Your_Sequence 8.0 -1.0 1 21 586 606
UGCAGUUGUUGUCUCAAGCUU ::::.: ..:.::.: AGAACAGAGAUACUGAUUGUA
Cleavage 8

ssp-miR444c-3p Your_Sequence 8.0 -1.0 1 21 7337 7357
UGCAGUUGUUGUCUCAAGCUU :: :::..:. :.: :: GGCCUACAGAUGAUCAUUUCA
Cleavage 8

ssp-miR396 Your_Sequence 8.25 -1.0 1 21 5368 5390 UUCCACAGCUUU--
CUUGAACUG : :::: .:::.:: :::: UCUUGCAAGCAGAAGUUGAGGAA Cleavage
2

ssp-miR156 Your_Sequence 8.5 -1.0 1 21 566 586
UGACAGAAGAGAGUGAGCACA : :: ::: ::::.: GGUAAACCCUAUCUCCUGUUA
Translation 3

ssp-miR159a Your_Sequence 8.5 -1.0 1 21 6609 6629
UUUGGAUUGAAGGGAGCUCUG :: :: ::: ..:: ::.: CAAAGGAACCUGUGAUGCAGA
Translation 4

ssp-miR166 Your_Sequence 8.5 -1.0 1 21 4485 4505
UCGGACCAGGCUUCAUUCCCC ::: ::::. ::..: GAAGAACAAAGCUGUGUUUGU
Cleavage 5

ssp-miR168a Your_Sequence 8.5 -1.0 1 21 3451 3471
UCGCUUGGUGCAGAUCGGGAC ::: :::::::. : AAUGAAUUCUUCACCAAGUUA
Translation 3

ssp-miR168a Your_Sequence 8.5 -1.0 1 21 4046 4066
UCGCUUGGUGCAGAUCGGGAC :: .::.:: ::: :. GGGAAGAAUUGUACAAAGAGG
Cleavage 3

ssp-miR396 Your_Sequence 8.5 -1.0 1 21 79 104 UUCCACAGCU-----
UUCUUGAACUG ::::.:::.: :::::: :: GAGUUUAAGGACAACUAGCUGUGCAA
Translation 2

ssp-miR437b Your_Sequence 8.5 -1.0 1 21 7632 7652
AAAGUUAGACAAGUUUGACAU : ::.. .::: .::.: : ACGUUGCCUUUGCUUAGCAUC
Cleavage 2

ssp-miR444b.1 Your_Sequence 8.5 -1.0 1 21 7458 7479 UUGUGG-
CUUUCUUGCAAGUUG :.:. :...:::: .:..:: CGAUCAGUGGGAAAGAUCGUAA Cleavage
16

ssp-miR444b.1 Your_Sequence 8.5 -1.0 1 21 4574 4594
UUGUGGCUUUCUUGCAAGUUG : ::: .::::: ..: CUUUAUACAAUGAAGCCUUGA
Translation 16

ssp-miR444b.1 Your_Sequence 8.5 -1.0 1 21 6550 6570
UUGUGGCUUUCUUGCAAGUUG : ::::::. .:: :. GCUGAUCCAAGAAGCUCAGAG
Cleavage 16

ssp-miR444b.1 Your_Sequence 8.5 -1.0 1 21 3568 3587
UUGUGGCUUUCUUGCAAGUUG .: :::: ::::.: ..:: UACCUUG-AAGAGAUAUGCAC
Cleavage 16

ssp-miR444b.1 Your_Sequence 8.5 -1.0 1 21 3262 3282
UUGUGGCUUUCUUGCAAGUUG ::: : .:::. ::.:: : CAAAUCAAGAGAGUGCUACUA
Cleavage 16

ssp-miR444b.1 Your_Sequence 8.5 -1.0 1 21 1360 1379
UUGUGGCUUUCUUGCAAGUUG .:: :: :::.::: .:. AGACCUG-AAGGAAGAUAUCU
Cleavage 16

ssp-miR444b.1 Your_Sequence 8.5 -1.0 1 21 5226 5245
UUGUGGCUUUCUUGCAAGUUG :: :: .:::: : .::: AUACAUGGGAGAA-GAUACAU
Cleavage 16

ssp-miR444c-3p Your_Sequence 8.5 -1.0 1 21 4381 4403
UGCAGUUGUUGUCU--CAAGCUU :: :: : :::::.: ..::: AAACUAGCCAGACAGCUGUUGCU
Cleavage 8

ssp-miR444c-3p Your_Sequence 8.5 -1.0 1 21 3235 3255
UGCAGUUGUUGUCUCAAGCUU ::. :::.::: ::. ACAAAUGGAACAGCAAAUGUU
Cleavage 8

ssp-miR444c-3p Your_Sequence 8.5 -1.0 1 21 2906 2925
UGCAGUUGUUGUCUCAAGCUU ::.:::::. :::: CUGGAUGGGACAAUU-CUGCA
Cleavage 8

ssp-miR528 Your_Sequence 8.5 -1.0 1 21 7584 7604
UGGAAGGGGCAUGCAGAGGAG : :: ::..::.:..:: AUGCUGUGUGUGUCUUUUGGC
Cleavage 1

ssp-miR827 Your_Sequence 8.5 -1.0 1 21 5743 5763
UUAGAUGACCAUCAGCAAACA :::.:: ::: ::: CCAAGACUGGUGUUCAACUAC
Cleavage 3

ssp-miR827 Your_Sequence 8.5 -1.0 1 21 592 611
UUAGAUGACCAUCAGCAAACA : ::::: :: ::::: GAGAUACUGAUUGU-AUCUAC
Translation 3
